# Protein Kinase A is a Functional Component of Focal Adhesions

**DOI:** 10.1101/2023.08.18.553932

**Authors:** Mingu Kang, Amanda J. Senatore, Hannah Naughton, Madeline McTigue, Rachel J. Beltman, Andrew A. Herppich, Mary Kay H. Pflum, Alan K. Howe

## Abstract

Focal adhesions (FAs) form the junction between extracellular matrix (ECM)-bound integrins and the actin cytoskeleton and also transmit signals that regulate cell adhesion, cytoskeletal dynamics, and cell migration. While many of these signals are rooted in reversible tyrosine phosphorylation, phosphorylation of FA proteins on Ser/Thr residues is far more abundant yet its mechanisms and consequences are far less understood. The cAMP-dependent protein kinase (protein kinase A; PKA) has important roles in cell adhesion and cell migration and is both an effector and regulator of integrin-mediated adhesion to the ECM. Importantly, subcellular localization plays a critically important role in specifying PKA function. Here, we show that PKA is present in isolated FA-cytoskeleton complexes and active within FAs in live cells. Furthermore, using kinase-catalyzed biotinylation of isolated FA-cytoskeleton complexes, we identify fifty-three high-stringency candidate PKA substrates within FAs. From this list, we validate tensin-3 (Tns3) – a well-established molecular scaffold, regulator of cell migration, and component of focal and fibrillar adhesions – as a novel direct substrate for PKA. These observations identify a new pathway for phospho-regulation of Tns3 and, importantly, establish a new and important niche for localized PKA signaling and thus provide a foundation for further investigation of the role of PKA in the regulation of FA dynamics and signaling.

## INTRODUCTION

Focal adhesions (FAs) are multi-molecular assemblages, consisting of hundreds of interconnecting proteins (1,2), that couple the cytoplasmic tails of extracellular matrix (ECM)-bound integrins to the actin cytoskeleton (3,4). FAs are also dynamic, and iteratively assemble, mature, and disassemble to providing strong yet reversible and regulatable adhesion. In addition to their role as dynamic physical couplings, FAs are *bona fide* centers for transmitting signals initiated by integrin engagement (5–9). Importantly, given their physical and functional position between the force-generating actin cytoskeleton and the force-resisting extracellular matrix, FAs are subject to mechanical inputs that regulate their assembly, conformation, composition, and disassembly. Such inputs have been widely demonstrated to alter the conformation of many structural proteins within FAs. At some point in this process, mechanically-instigated changes in protein conformation are ‘converted’ into changes in enzymatic activity such that mechano*sensation* gives rise to mechano*transduction* (10). Thus, FAs receive and transmit information about both the biochemical *and* the mechanical nature of cell adhesion, and the complement of signals generated by FAs not only feedback to control cell adhesion & cytoskeletal dynamics, but also expand to regulate cell proliferation, survival, and differentiation (6,8). Elucidation of the interactions and activities of various structural and signaling components within FAs will provide deeper understanding of how these structures are regulated and, in turn, how they regulate other cellular behaviors.

An important mechanism through which FAs integrate their structure, their remodeling, and their signaling is through reversible phosphorylation of most if not all FA components (5,11). Of note, FAs are especially enriched in phosphotyrosine (pTyr), a modification that represents less than 2% of *total cellular* protein phosphorylation but ∼25% of the phosphorylation found in isolated focal adhesion/cytoskeleton (FACS) fractions (11). While the importance of phosphotyrosine to FA function is central and undeniable, it is equally undeniable that serine/threonine phosphorylation [pSer/pThr] is far more abundant in FA than pTyr (11). Also, examination of published literature (11) and of publicly-available protein phosphorylation data repositories (*e.g.* PhosphoSitePlus, Phosida, dbPAF) shows that essentially every FA protein known to contain pTyr also contains pSer and/or pThr (*unpublished observations*). However, despite its prominence in regulating protein and cell function and its prevalence in FAs, the role of Ser/Thr phosphorylation in the regulation of FAs and their components is comparatively – perhaps *vastly* – understudied. A more complete understanding of how FAs are regulated and how they engage signaling cascades must therefore include direct and thorough characterization of the mediators, targets, and consequences of Ser/Thr phosphorylation within FAs.

A promising candidate in this regard is the cAMP-dependent protein kinase, PKA. PKA is a ubiquitous and promiscuous kinase with hundreds of substrates across the cellular landscape. PKA is a heterotetramer of a regulatory (R) subunit dimer associated with two catalytic (C) subunits. In response to a wide variety of stimuli, cAMP increases and binds R subunits, promoting the activation of C subunits (12). The classical model of PKA activation involves dissociation & diffusion of C subunits from cAMP-bound R subunits. However, impactful recent work suggests that the catalytically active form of PKA remains an intact, tetrameric holoenzyme (13), suggesting that PKA may act in a far more localized or spatially-restricted manner than previously thought. PKA activity is further parsed into a large and growing number of physically and functionally discrete signaling nodes, organized by a diverse set of scaffolds known as AKAPs (A-kinase anchoring proteins (14–16)). Each AKAP has a distinct complement of additional domains that mediate interaction with other proteins and thus direct the distinct targeting and function of various AKAP-PKA complexes within cells (14–16).

PKA is both a complex effector and regulator of cell migration (17–19), with the balance of its effects achieved through localization to distinct subcellular regions and structures associated with motility, such as the leading edge (20–24), the trailing edge (25,26), and the actin cytoskeleton (27,28). Thus, a fuller understanding of PKA-mediated regulation of migration requires a better understanding of how PKA activity is specified within motility-associated regions or structures. Importantly, PKA has been shown to be both activated and inhibited upon integrin engagement and adhesion to ECM (21,29–36), though the mechanistic details of this regulation are completely unknown. Furthermore, activation of PKA has been shown to either enhance or impede cell adhesion (17,30,36–42), in a cell- and/or ECM-specific manner. Finally, FAs contain a number of *bona fide* PKA substrates (17), including Ena/VASP proteins (43), the LIM & SH3 domain protein (LASP-1 (44)), Src-family kinases (Src (45,46) and Fyn (41)), SHP2/PTPN11 (47,48), and integrin α4 (49). To this end, we set out to determine if PKA is a functional constituent of FAs.

## RESULTS

### Inhibition of PKA affects FA morphology, distribution, and dynamics

Given that PKA has been shown to be activated upon integrin engagement and adhesion to ECM and that PKA activity has been shown to both enhance and impede cell adhesion in various cell types (17,19), we assessed the effect of inhibition of PKA on FA morphology and distribution during fibroblast cell adhesion and spreading. REF52 (a rat embryonic fibroblast cell line) were plated onto fibronectin (FN)-coated coverslips for varying amounts of time in the presence or absence of Rp-8-CPT-cAMPS (‘Rp-cAMPS’), a cAMP analogue that specifically and competitively binds PKA R subunits and prevents holoenzyme dissociation and kinase activation. Cells were then fixed and stained with antibodies against vinculin, talin, or paxillin to visualize FA. While there was no discernable difference in cell size or morphology between untreated and inhibitor-treated cells plated on FN, there were striking differences in FA morphology and distribution (**Fig. 1**). Specifically, at 1 h after plating, vinculin-containing adhesion complexes in PKA-inhibited cells were larger and more peripherally located (**Fig. 1A**). Visualization and quantification of the skewed peripheral distribution was facilitated by conversion of round cell images to linear images using polar transformation (**Fig. 1, A and B**). The largest adhesion complexes, reminiscent of clustered or aggregated complexes seen upon inhibition of longitudinal FA splitting (50), precluded an accurate determination and comparison of individual FA size and morphometrics. We therefore assessed FA morphology at later time points after plating. At 1.5, 2, 3, and 4 h, paxillin- or talin-positive FAs in both populations were more discernable, but were significantly larger, longer, and more skewed to the periphery in PKA-inhibited cells compared to untreated cells (**Fig. 1, C-I; Fig. S1**). We also investigated the effects of PKA inhibition on PKA dynamics by live-cell imaging of REF52 cells expressing mCherry-tagged paxillin and plated onto FN-coated imaging dishes. Control-treated cells exhibited characteristic FA dynamics, as visualized by iterative formation and turnover of FA 30-90 min after plating (**Fig. 1F**), while PKA inhibitor-treated cells exhibited larger, more peripheral, and far less dynamic FAs (**Fig. 1F**), reminiscent of the adhesions observed in fixed cell experiments (**Fig. 1, A-E**; **Fig. S1**). Of note, global activation of PKA by addition of forskolin (*Fsk*) and isobuylmethylxanthine (*IBMX*), which elevate total intracellular cAMP, inhibited cell spreading and FA formation (**Fig. S2**), consistent with earlier observations (reviewed in (17,19)). Taken together, these observations demonstrate that regulated PKA activity is required for normal FA morphology and dynamics.

**Figure 1.**
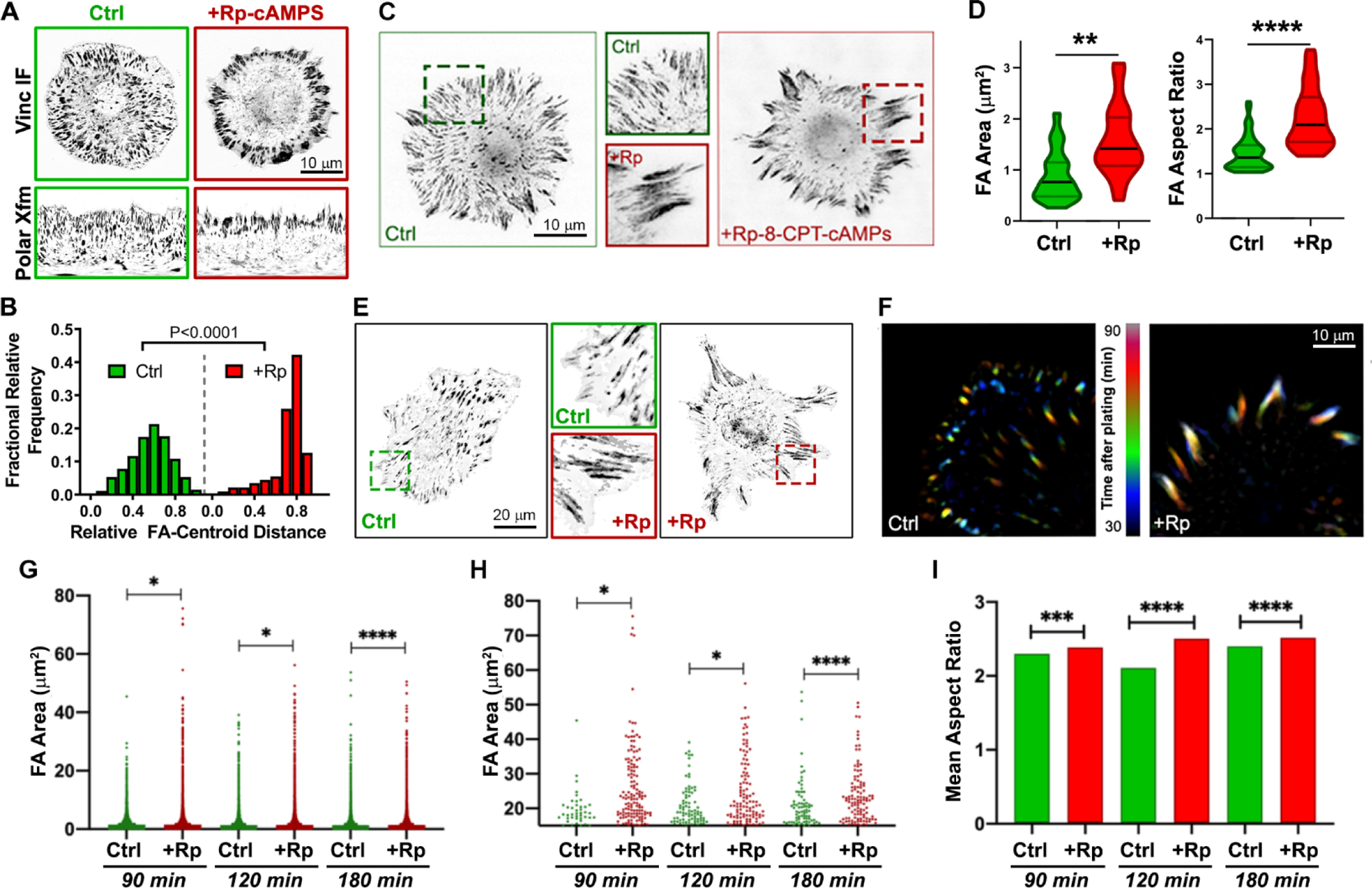
Inhibition of PKA alters FA morphology, distribution, and dynamics in spreading cells. (**A**) Anti-vinculin immunofluorescence of REF52 cells plated onto fibronectin (FN)-coated coverslips for 1 h in the absence (*Ctrl*) or presence of an inhibitor of PKA activity (Rp-8-CPT-cAMPS (*+Rp-cAMPS*); 50µM). Images are shown using an inverted, grayscale lookup table for ease of visualization. To facilitate analysis of FA distance from the cell center, images were ‘unwrapped’ using a clockwise polar transformation (*Polar Xfm*; bottom panels). (**B**) Histograms of relative FA distance from cell centroid (0 = cell center; 1 = cell periphery) in control- and PKA inhibitor-treated cells (600 and 601 FAs for Ctrl and +Rp-cAMPS samples, respectively). Distributions were analyzed using a Kolmogorov-Smirnov test (K-S distance = 0.5058). (**C**) Anti-paxillin immunofluorescence of REF52 cells 1 h after plating on FN in the absence (*Ctrl*) or presence of Rp-8-CPT-cAMPS. Enlargements of the dash-outlined areas in each panel are shown in the middle column. (**D**) FA morphometrics from control- and PKA inhibitor-treated cells. Violin plots summarize data from 3 experiments (2784 and 2155 total FAs for Ctrl and PKA-inhibited samples, respectively), with the width of the plot proportional to the number of points, top and bottom depicting the maximum and minimum values, and black lines depicting the median and colored lines depicting the 25^th^ and 75^th^ quartiles (**, p<0.05; ****, p<0.005). (**E**) Anti-talin immunofluorescence of REF52 cells plated onto FN for 4h in the absence or presence of PKA inhibitor. The middle column shows enlargements of the dash-outlined areas in each panel. (**F**) Temporal color-coded live-cell images of mCherry-paxillin-expressing REF52 cells spreading on FN for 1 h min the absence or presence of Rp-8-CPT-cAMPS. Image acquisition began 30 min after plating. (**G-I**) Immunofluorescence images were analyzed at 90, 120 and 180 minutes to determine FA size (**G, H**) and aspect ratio (**I**). (**G**) Depicts all measured adhesion values at each timepoint. (**H**) Depicts all measured adhesion values greater than 15 μm^2^ to highlight differences at larger adhesion area values. (**I**) Depicts mean aspect ratio for all groups, analyzed by unpaired t-tests (n = >170 cells from 2 experiments; *p<.05, *** p<.001, **** p<.0001).

### PKA is present in isolated FA-cytoskeleton preparations

Although the aforementioned results support a requirement for PKA activity in FA regulation, the effect of the globally-applied PKA inhibitor may well be mediated through various mechanisms acting in multiple subcellular compartments. However, given that a number of *bona fide* PKA substrates reside within FAs (17,19,51), we hypothesized that FAs might contain a specific, discrete pool of PKA. To investigate this, we modified a published method (52) for isolating cellular fractions that are highly enriched for FA/cytoskeleton (FACS) components. Briefly, cells are plated onto ECM-coated dishes, treated with a cleavable and membrane-permeable cross-linker, unroofed by detergent lysis and fluid shear, and the remaining FACS proteins are analyzed *in situ* by fluorescence microscopy or recovered from the surface for immunoblotting (see ***Materials and Methods*** for details). Immunofluorescence and immunoblotting analyses of FACS preparations confirmed the removal of the majority of cellular material and retention and enrichment of FA proteins and FA-associated F-actin (**Fig. 2, A-D**). Importantly, PKA RII subunits were also retained in these FACS preparations, as assessed by immunofluorescence or immunoblotting (**Fig. 2, B and C**) and while some of the PKA signal seemed to be associated with residual, presumably ECM-proximal membrane fragments, as assessed by fluorescence microscopy, a significant fraction co-localized with actin-associated FAs (**Fig. 2B**). The PKA catalytic subunit was also found within purified FACS preparations (**Fig. 2D**). These observations confirm earlier studies in which PKA subunits were found in unbiased/shotgun proteomics analyses of adhesion complexes isolated by various means (1,51,53) and demonstrate that a pool of PKA resides within FAs.

**Figure 2.**
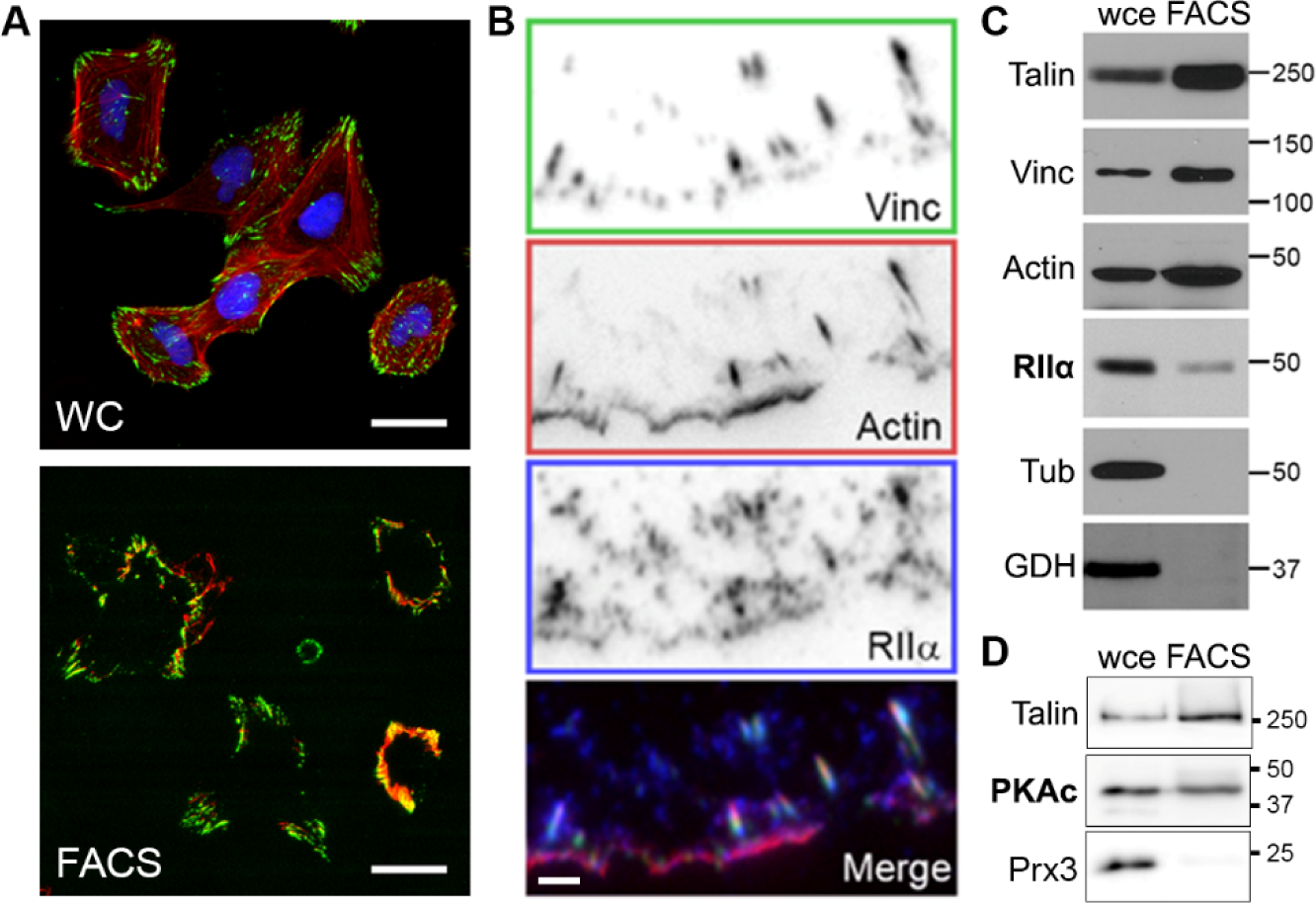
PKA type-II regulatory (RII) subunits are present in isolated focal adhesion/cytoskeleton (FACS) fractions. **(A**) REF52 cells plated on FN-coated coverslips were fixed with DSP without (for whole cells, *top*) or with (for FACS preparation, *bottom*) subsequent fluid shear, then stained for talin (*green*), F-actin (*red*), and nuclei (*blue*). Scale bar = 30 µm. (**B**) REF52 cells were FACS-prepped then stained to visualize vinculin (*green*), F-actin (*red*), and PKA RIIα subunits (*blue*). Scale bar = 5 µm. (**C, D**) Ten micrograms of whole-cell extract (*wce*) or isolated FACS proteins (*FACS*) were separated by SDS-PAGE and immunoblotted with antibodies against talin, vinculin (*Vinc*), actin, PKA RIIα (*RII*α) or catalytic (*PKAc*) subunits, tubulin (*Tub*), GAPDH (*GDH*), and peroxiredoxin-3 (*Prx3*). Molecular weight markers (in kDa) are shown on the right.

### Phospho-RII subunits, autophosphorylated at a site associated with PKA activation, localize to focal adhesions

The presence of PKA RII subunits in FAs suggests that PKA might be active within these structures. To address this, we took advantage of recent work establishing that an epitope on the inhibitory loop of RII subunits containing an autophosphorylation site (Ser99) is exposed upon cAMP-induced dissociation of the PKA C subunit (54). This phosphorylation of RII subunits precedes cAMP binding, but cAMP-binding and activation of PKA displaces the C subunit, allowing increased access of a phospho-specific antibody to this epitope; thus, detection of the phospho-RII epitope can be used in immunofluorescence as an indicator of highly-localized PKA activity within anchored holoenzyme complexes (54). Fixation and unroofing of REF52 cells followed by immunostaining with antibodies against vinculin and against phospho-Ser99 RII subunits (*pSer99*), revealed a striking colocalization of phospho-Ser99 RII within FAs (**Fig. 3A**). Importantly, phospho-Ser99 RII signal was essentially eliminated by phosphatase treatment of the samples prior to staining, confirming the phosphorylation-specificity of the immuno-reactivity (**Fig. 3A**). The prominent colocalization of phospho-Ser99 with vinculin, as well as its decrease upon phosphatase treatment, was confirmed through quantification of colocalization using two distinct methods - Pearson’s correlation coefficients, which determine pixel-by-pixel covariance in two images independently of signal intensity (**Fig. 3B**) and Li’s intensity correlation coefficients (ICQ), which assess covariance of signal intensities in two fluorescence channels (**Fig. 3C**).

**Figure 3.**
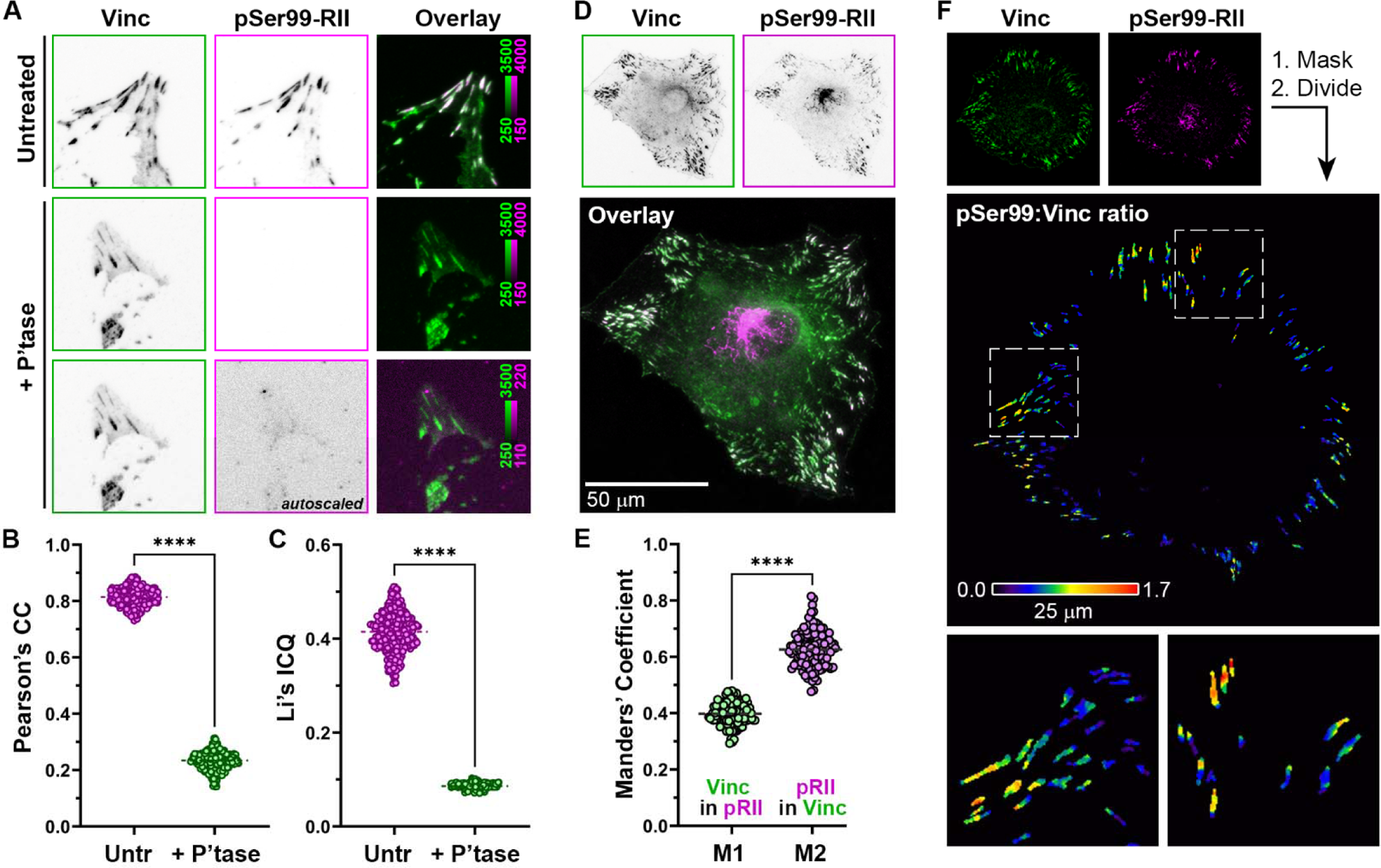
Phosphorylated RII subunits PKA localize to FAs. (**A**) FN-adherent REF52 cells were fixed, unroofed, left untreated or incubated with lambda phosphatase (+ *P’tase*), then immunostained with antibodies against vinculin (*Vinc*) and against autophosphorylated PKA RII subunits (*pSer99-RII*). (**B**, **C**) Colocalization of pSer99-RII and vinculin with and without phosphatase treatment was analyzed using Pearson’s correlation coefficient (*CC*) and Li’s intensity correlation quotient (*ICQ*). Data represent 300 100 x 100 µm microscopic fields from two different experiments and were analyzed by unpaired t-test (****, p<0.001). (**D**) FN-adherent REF52 cells were fixed and immunostained (without unroofing) with antibodies against vinculin (*Vinc*) and against autophosphorylated PKA RII subunits (*pSer99-RII*). (**E**) Manders’ overlap coefficients were calculated to determine relative amount of vinculin signal that overlaps with pSer99-RII (*M1*) and of pSer99-RII that overlaps with vinculin (*M2*). Data represent 120 cells from in 3 experiments. (**F**) REF52 cells were fixed and stained as described for panel (**D**). Vinculin images were used to generate binary mask images that were multiplied input vinculin and pSer99-RII images to eliminate non-FA signal. Ratiometric images were generated by dividing pixel-by-pixel pSer99-RII intensities by corresponding vinculin intensities and pseudocolored using the S-Pet lookup table in Fiji.

Given the rather striking abundance of phospho-Ser99 RII in unroofed FAs, we investigated whether this localization could also be visualized in intact (*i.e.* non-unroofed) cells. Immunofluorescence of non-unroofed cells showed strong pSer99-RII staining in the Golgi as well as prominent localization to FAs (**Fig. 3D**). Cursory inspection of such images showed that some vinculin-containing FAs showed little-to-no phospho-Ser99 RII signal (**Fig. 3D**) and calculation of Mander’s colocalization coefficients confirmed that the proportion of phospho-Ser99 colocalized with vinculin is higher than the proportion of vinculin colocalized with phospho-Ser99 (**Fig. 3E**). These data suggest that phospho-Ser99 RII subunit intensity – and thus PKA activity – is higher in some FAs than others, suggesting some degree of sub-cellular and/or functional heterogeneity or specificity. This was confirmed by ratiometric analysis of phospho-RII and vinculin staining intensities within FAs which also revealed that the relative abundance of phospho-Ser99 RII was not uniform/homogeneously distributed within individual FAs, but rather localized to sub-adhesion ‘hot-spots’ (**Fig. 3F**). These observations demonstrate that a phospho-epitope in RII subunits that is exposed upon PKA activation is a prominent feature of FAs.

### PKA is dynamically regulated within focal adhesions

Prolonged exposure of the RII Ser99 epitope increases the likelihood of its recognition by phosphatases, leading to dephosphorylation and loss of immunoreactivity; thus, immuno-reactivity with phospho-Ser99 RII antibody is taken to indicate a pool of PKA activity that is hyper-localized and/or highly dynamic (54). The presence and non-uniform distribution of phospho-Ser99 RII subunits in FAs suggests that PKA might be active and dynamically regulated within these structures. To address this, we generated a <u>pax</u>illin-linked ratiometric <u>A</u>-<u>k</u>inase <u>s</u>ensor (PaxRAKS), a focal adhesion targeted biosensor that allows us to monitor PKA activity within adhesive structures (**Fig. 4, A and B**). For the activity sensing module, we used GAkdY (55), a single-fluorophore PKA biosensor module in which the fluorescence intensity of a conformationally-sensitive GFP is controlled by a PKA activity sensing domain derived from A-kinase activity reporter 2 (AKAR2 (56)), comprising a phosphothreonine binding domain linked to a RRAT (single amino acid code) PKA substrate sequence. This module was fused to the C-terminal end of the FA adaptor protein paxillin to drive targeting of the sensor to adhesion complexes. Finally, the red fluorescent protein mCherry was fused to the N-terminus of paxillin-GAkdY to serve as an internal control for fluctuations in biosensor abundance during FA assembly and turnover. This provides a ‘denominator’ for PKA-mediated changes in GFP intensity such that the ratio of GFP to mCherry intensity reflects changes in PKA activity within FA.

**Figure 4.**
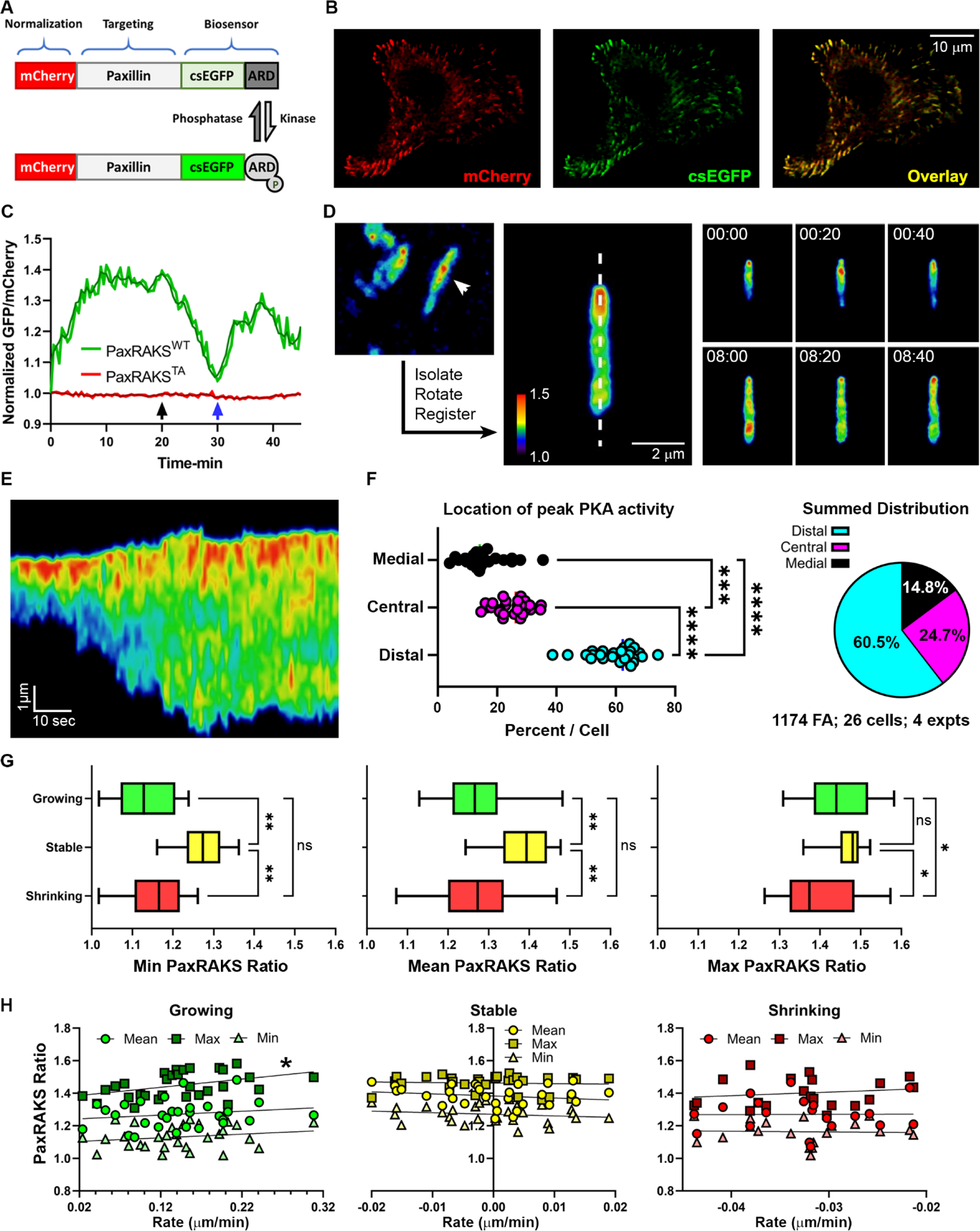
A FA-targeted PKA biosensor, PaxRAKS, reveals spatially and temporally dynamic PKA signaling events within individual FAs. (**A**) Schematic of PaxRAKS (*Pax*illin-fused *R*atiometric *A*-*K*inase *S*ensor). A PKA activity-responsive domain (*ARD*) coupled to a conformation-sensitive EGFP (*csEGFP*) is fused to the C-terminus of paxillin for localization to FAs. The monomeric red fluorescent protein mCherry is fused to the paxillin N-terminus as a denominator, allowing normalization of phosphorylation-dependent changes in EGFP fluorescence to the local abundance of the biosensor. (**B**) Images of a PaxRAKS-expressing cell, depicting simultaneous capture and spatial coincidence of the PKA-responsive csEGFP and reference mCherry signals. (**C**) Ratios of csEGFP:mCherry intensity over time in selected, individual FAs containing either wild-type PaxRAKS (*WT*; *green*) or a phospho-resistant Thr/Ala point mutant (*TA; red*). Black and blue arrows indicate the addition and wash-out, respectively, of Rp-8-CPT-cAMPS. Light and dark lines depict raw ratio values (acquired at 3 frames/min) and boxcar-averaged (−/+ 1 time point) values for smoothing, respectively. (**D**) Color-coded Ratiometric (csEGFP:mCherry) PaxRAKS signal in individual FAs, color-coded using a rainbow lookup table, such that low ratios, indicating low PKA activity, are depicted in cooler colors and high ratios (*i.e.* high activity) in warmer colors (*left*; see Supp. Movie 1). A set of ImageJ macros was used to isolate, vertically align, and spatially register individual adhesions (*middle*) to aid in visualization (*right*) and subsequent analysis (*see below;* Supp. Movie 2). Time-stamped panels show the localization and dynamics PKA activity within an individual FA at the indicated time points (*min:sec*). (**E**) Linescan analysis of PaxRAKS dynamics over 90 sec under the region of interest depicted by the dotted line in the FA depicted in panel (**C**). Signal is plotted along the y-axis as a function of time, along the x-axis. (**F**) Analysis of location of peak PKA activity (*i.e.* maximum PaxRAKS ratio value) in 300 FA from 26 cells in 3 experiments. For each cell, 8-12 FA were imaged every 3 sec for 1 min, kymographs were produced as described for panel (**E**), and the peak signal was scored as being located in the distal, central, or medial (relative to the center of the cell) third of the trace. Locations were tallied on a per-cell basis. Data are presented for each cell (scatter graph, *left*; ***, p<0.005; ****, p<0.001). and summarized for all cells (pie chart, *right*). (**G**) Minimum, mean, and maximum PaxRAKS ratios in growing, stable, and shrinking adhesions. Peripheral FAs with lengths that changed at rates above +0.02 µm/min for at least 5 min were considered *Growing* (n=30), those with rates between +0.02 and −0.02 were considered *Stable* (n=30), and those with rates below −0.02 µm/min considered *Shrinking* (n=25). Data were analyzed using ordinary one-way ANOVA with multiple comparisons (*, p<0.05; **, p<0.01; *ns*, not significant). (**H**) The same data depicted in (**G**) was re-plotted as XY scatter plots of minimum, maximum, and mean PaxRAKS ratio values *versus* growth rate magnitude in individual FAs. The only significant ratio/rate correlation was between maximum ratio and growth rate in growing FAs (*, r^2^ = 0.174, p =0.0217).

Live-cell imaging of cells transfected with PaxRAKS confirmed proper localization of the biosensor to FA (**Fig. 4B**). To confirm the specificity of PaxRAKS, we calculated the GFP:mCherry ratio of FA in cells before and after treatment with a specific PKA inhibitor (Rp-cAMPS) as well as after inhibitor washout. The ratio showed some minor fluctuations and a significant increase over time (**Fig. 4C**, *green line*). Importantly, the ratio dropped rapidly upon inhibition of PKA (**Fig. 4C**, *black arrow*) and recovered rapidly after inhibitor washout (**Fig. 4C**, *blue arrow*). Furthermore, a phospho-resistant mutant of PaxRAKS, in which the Thr phospho-acceptor residue is mutated to Ala (PaxRAKS^TA^), showed lower fluctuations and peaks in GFP:mCherry ratio in untreated cells and no changes in response to modulation of PKA activity (**Fig. 4C**, *red line*). Taken together, these observations support the conclusion that PaxRAKS is a functional and specific biosensor for detecting PKA activity within FA.

Preliminary analyses revealed patterns of considerable variety, complexity and dynamicity of PKA activity within individual FA in migrating cells (**Fig. 4, D and E; Movie S1 and Movie S2**). However, a few characteristics were particularly obvious and noteworthy. First, PKA activity within adhesions was highly dynamic and fluctuated rapidly during both FA growth and disassembly (**Fig. 4, D and E; Movie S2**). Second, PKA activity was not distributed evenly throughout FA; rather, a significant percentage of FA exhibited peaks of PKA activity in the distal (*i.e.* towards the cell periphery) third of their area, with peaks occurring in the central and medial thirds with lower frequency (**Fig. 4, D-F**). Correlating these patterns of PKA activity with aspects FA dynamics could lend significant insight into the function of this pool of PKA. However, the parameter space of dynamic FAs is large, consisting of size, aspect ratio, assembly/disassembly rates, and position within the cell – all of which requires sophisticated image analysis and tracking software and makes this analysis complex and worthy of concerted, rigorous effort. Nonetheless, to begin to assess whether there are any correlations between the dynamics of FAs and the PKA activity within them, we determined the minimum, maximum, and mean PaxRAKS ratio values in individual FAs. We chose FAs near the cell periphery, as those tend to be the most dynamic, and assigned FAs as ‘growing’ (with lengths that changed at rates > +0.02 µm/min for at least 5 min), ‘stable’ (with rates between +0.02 and −0.02 µm/min for 5 min) and ‘shrinking’ (with rates < −0.02 µm/min for 5 min). Of note, the dataset and analyses here include FAs that exhibited one behavior then transitioned into another (*e.g.* values within an FA that assembled for ≥ 5 min but then stabilized for ≥ 5 min were included in both ‘growing’ and ‘stable’ analyses, respectively). Comparison of minimum, mean, and maximum PaxRAKS ratios between growing, stable, and shrinking FAs revealed that stable peripheral FAs showed higher minimum and average PaxRAKS ratios (*i.e.* PKA activity) compared to assembling or disassembling FAs (**Fig. 4G**). Also, peak PaxRAKS values were comparable between growing and stable adhesions but slightly lower in shrinking FAs (**Fig. 4G**). Re-analysis of minimum, maximum, and mean PaxRAKS ratio values as a function of the *magnitude* of growth rate, as XY scatter plots, revealed no strong correlation between PKA activity and the magnitude of FA assembly or disassembly rate, save for a slight but statistically significant (r^2^ = 0.174, p =0.0217) correlation between maximum ratio and growth rate in growing FAs (**Fig. 4H**). These preliminary analyses suggest that the dynamics of PKA activity within FAs seem to vary with the state, rather than the rate, of FA growth or disassembly and underscore the importance of further investigation of how these patterns correlate with FA dynamics and location. Nonetheless, the current observations, along with those presented in **Fig. 2**, clearly establish that PKA is present and active within FA and exhibits dynamic fluctuations in activity with sub-FA spatial localization.

### Kinase-catalyzed biotinylation of FACS preparations identifies novel candidate FA PKA substrates

The fact that PKA is present and active in FA suggests that FA likely contain PKA substrates, an hypothesis supported by numerous prior reports reviewed elsewhere (17). To identify novel FA substrates for PKA, we used kinase-catalyzed biotinylation (57) where lysates from FACS preparation were incubated with an ATP analog bearing biotin at the γ-phosphoryl position (ATP-biotin) and purified PKA catalytic subunit (PKA-C), which transfers the terminal phosphorylbiotin from ATP-biotin to PKA substrates. The biotinylated substrate proteins can then be recovered by streptavidin enrichment and identified by liquid chromatography-tandem mass spectrometry (LC-MS/MS). We first performed proof-of-principle biotinylation experiments to determine whether ATP-biotin labeling could be successfully applied to FACS. Typical kinase-catalyzed biotinylation reactions used 200-1000 µg of lysate per reaction (57). However, the highly selective nature of the FACS preparation, which typically yielded 5-10 µg per 177 cm^2^ (15 cm diameter) culture dish, made it infeasible to iteratively and reliably generate this amount. Thus, a second goal was to assess if robust biotinylation would be observed with more limited amounts of cell lysate. FACS preparations were incubated with ATP-biotin both without and with recombinant PKA-C before separation by SDS-PAGE and visualization of biotinylated protein using streptavidin-conjugated horseradish peroxidase. A low level of biotinylated proteins was observed without added PKA-C (**Fig. S3**, lane 2), which is consistent with the presence of active kinase activity in the preparation. Because ATP-biotin is compatible with all cellular kinases (58), addition of PKA-C to the ATP-biotin reaction allowed PKA substrates to be distinguished from substrates of other cellular kinases. In fact, the level of protein biotinylation was significantly elevated in the presence of PKA-C (**Fig. S3**, lanes 3 and 4), which is consistent with the presence of PKA substrates in the preparations and ideal for proteomics analysis. In addition, robust biotinylation was observed with as little as 5 µg of FACS lysate, a ten-fold lower amount than typically used previously (57). These biotinylation assays establish the feasibility of kinase-catalyzed biotinylation for identification of putative substrates from subcellular fractions.

After observing robust biotinylation by PKA-C in FACS preparations, subsequent preparative-scale reactions were performed. To identify FACS proteins modified by PKA-catalyzed biotinylation and, therefore, putative PKA substrates, using LC-MS/MS analysis. Four independent biotinylation experiments were performed using 30-40 µg of FACS lysate, each comprising three preparative-scale reactions in which FACS lysate was incubated with: (i) PKA with ATP (unlabeled control); (ii) ATP-biotin without PKA-C (negative control); (iii) ATP-biotin with PKA-C (experimental sample). Biotinylated products were enriched with streptavidin-conjugated beads and analyzed by LC-MS/MS. Quantitative proteomic analysis (59) identified a total of 730 proteins among the four trials (**Table S1A**). Proteins that were enriched in or exclusive to the unlabeled control samples (*i.e.* PKA-C, +ATP; sample 1) were removed from the list. Then, enrichment values were calculated by dividing the experimental sample (*i.e.* ATP-biotin, +PKA-C; sample 3) by the negative control (*i.e.* ATP-biotin, -PKA-C; sample 2), and two ‘hit lists’ were generated: one of 98 proteins with enrichment values greater than or equal to 1.5-fold in at least 2 of the 4 trials (**Table S1B**), which represents all possible hits, and a second, higher-stringency list of 53 proteins that were enriched greater than or equal to 6.0-fold in at least 2 of the 4 trials (**Table S1C**). Bioinformatic analysis using gene ontology (GO) showed biological processes (GOBP) with immediate relevance to cell adhesion, migration, and cytoskeletal dynamics as the most highly represented classes among the identified proteins (**Fig. 5A; Table S1C**), consistent with FA function. In addition, interactome analysis of the high-stringency hit list revealed significant physical interactions between and among PKA and the potential substrates (**Fig. 5B**), with 36 proteins either directly or indirectly interacting with PKA (69%). While the list included some known PKA substrates, such as ezrin (60) and galectins (61), most hit proteins had not been previously described as PKA substrates. However, analysis of the 53 high-stringency hits for possible PKA sites identified that 51 proteins (96%) contain one or more moderate-to-strong substrate motifs for PKA phosphorylation (**Fig. 5C**). These results provide a number of putative, novel PKA substrates that function in or on the cell adhesion and migration machinery.

**Figure 5.**
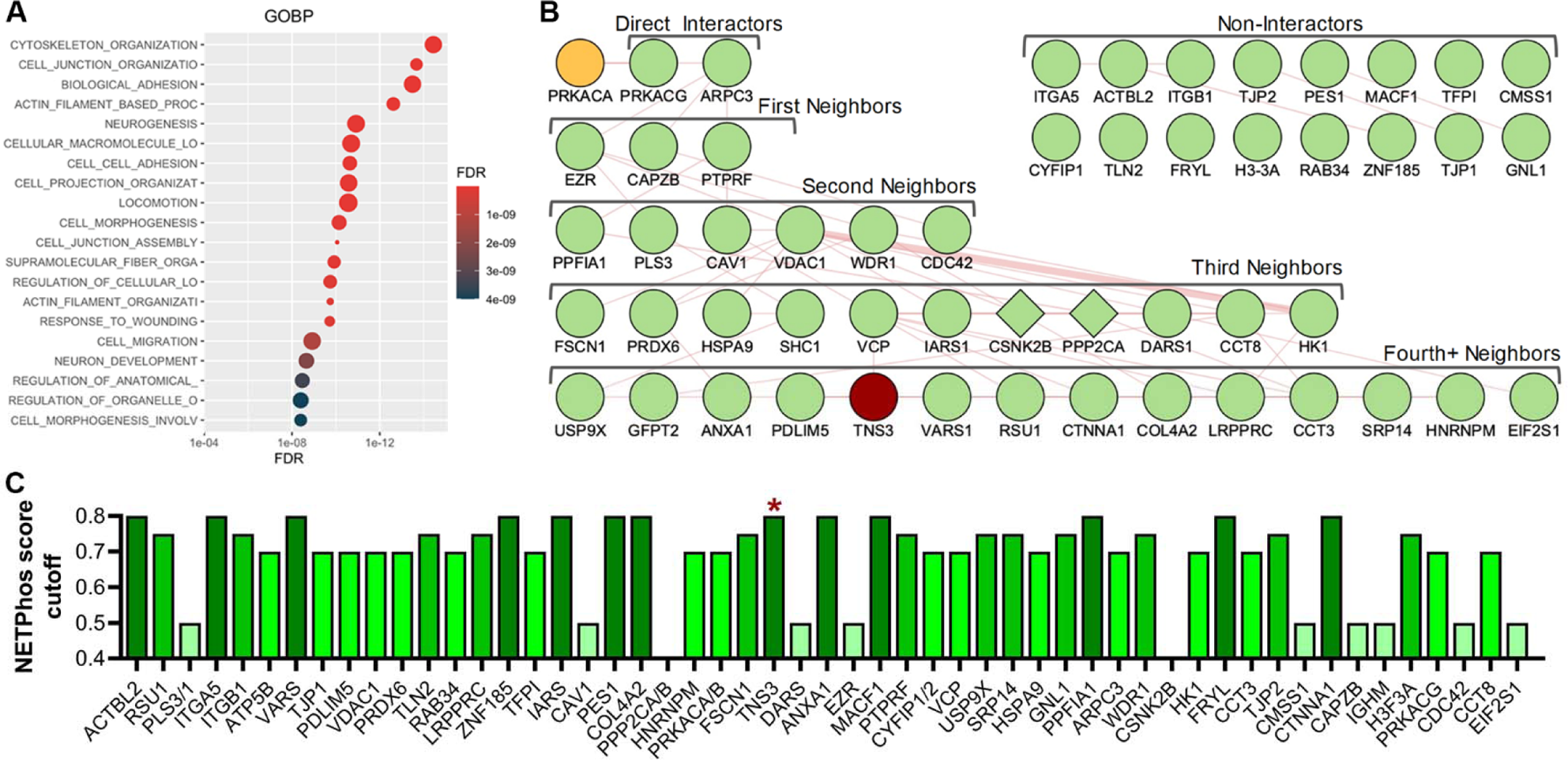
Candidate PKA substrates from kinase-catalyzed biotinylation of isolated focal adhesion/cytoskeleton fractions. Focal adhesion/cytoskeletal (FACS) proteins were isolated from REF52 cells stably adhered to FN-coated plates and incubated with ATP-biotin with or without exogenous PKA catalytic subunit. Biotinylated proteins were captured on immobilized avidin resin and processed for proteomic analysis by LC-MS/MS. (**A)** Analysis of PKA-catalyzed biotinylated FACS proteins by GOBP (Gene Ontology Biological Process). The 53 high stringency hit list of proteins (**Supp. Table S1C**) was analyzed for inclusion in the indicated GOBP classes. The size of the class circles is proportional to the number of hits in that class, while the shading represents the false discovery rate (FDR) as indicated in the scale. (**B)** The 53 high stringency hits (**Supp. Table S1C**) were analyzed using the GeneMANIA plugin in Cytoscape to identify and visualize known physical interactions between and among PKA and potential FACS substrates. The PKA catalytic subunit (*top, yellow*) is shown with direct interactors (*first row*) and indirect interactors separated by one degree (first neighbors, *second row*), two degrees (second neighbors, *third row*), or three or more degrees of freedom (third and fourth+ neighbors. *bottom two rows*). Proteins with no prior physical interactions with PKA are shown (non-interactors, two rows in the top right). The two proteins that did not contain PKA substrate sites (*see below*) are indicated as diamonds, while Tns3, which was chosen for further validation, is shaded in dark red. (**C)** Scoring the presence of putative PKA motifs in candidate substrates. The amino acid sequences of the high stringency hits were analyzed using NETPhos 3.1 to identify predicted PKA-specific phosphorylation sites. For each phosphoacceptor residue, this analysis generates a score representing the probability (from 0-1) that the residue is a PKA phosphosite based on the homology the flanking amino acids to known PKA consensus sequences. The bars indicate that the given protein contains at least one site with a score greater than the indicated cutoff of 0.5 (the NETPhos minimum to include only positive predicted sites), 0.7, 0.75, or 0.8. The red asterisk indicates the high cutoff score for Tns3.

### The FA protein Tensin-3 is a novel PKA substrate

From the list of 53 potential FA substrates for PKA, tensin-3 was chosen for further biochemical validation. Tensins are a family of FA-associated proteins with important roles in regulating cell adhesion, migration, and associated signaling events (62–64). Tensin-3 (Tns3) is a prominent component of FA but is also enriched in and necessary for fibrillar adhesions, elongated cell-ECM junctions that mature and centralize from more peripheral FA (62,65). Tns3 contains an active protein tyrosine phosphatase (PTP) domain as well as phosphotyrosine-binding SH2 and PTB domains and scaffolds a variety of motility-associated signaling molecules, including β1 integrins, FAK, p130Cas, PEAK1, DLC1, DOCK5 (62–64,66). Importantly, Tns3 expression is variably up- or down-regulated in various cancers and has often been demonstrated to suppress or govern cell motility and invasion (62–64,67–69). Finally, as with all Tns family members, Tns3 is extensively modified and regulated by phosphorylation (62–64,70,71) and while it contains several strong possible PKA substrate motifs (**Fig. 5C**), no prior connection between Tns3 and PKA has been reported. Thus, to corroborate the direct PKA-mediated modification of Tns3 revealed by *in vitro* biotinylation, we performed *in vitro* kinase assays by pulling down GFP-tagged Tns3 expressed in HEK293 cells and incubation with purified PKA catalytic subunit. Importantly, over-expressed GFP-Tns3 localized predominantly to FAs and more central fibrillar adhesions (**Fig. S4**). GFP-Tns3, but not GFP alone, was direct phosphorylated by PKA, as demonstrated by immunoblotting of reaction products with phospho-PKA substrate antibody (**Fig. 6A**). To determine whether Tns3 can be phosphorylated by PKA *in vivo*, we isolated GFP-tagged Tns3 expressed in HEK293 cells that were treated with Fsk and IBMX or DMSO and analyzed it by immunoblotting with a phospho-PKA substrate antibody. As expected, treatment with Fsk and IBMX activated PKA, as evidenced by increased reactivity of whole cell lysates with the phospho-PKA substrate antibody compared to control-treated cell lysates (**Fig. 6, B and C**). Importantly, GFP-Tns3 isolated from Fsk/IBMX-treated cells also showed increased anti-phospho-PKA substrate reactivity compared to control (**Fig. 6, B and C**), supporting the contention that Tns3 is phosphorylated by PKA *in vivo*. This is further supported by the observation that baseline level of phospho-Tns3 is decreased in cells treated with H89, a kinase inhibitor active against PKA (**Fig. 6C**). To determine whether endogenous Tns3 behaved comparably to exogenous GFP-Tns3, we utilized U2OS osteosarcoma cells, which express high levels of endogenous Tns3 (https://www.proteinatlas.org/). Tns3 immunoprecipitated from U2OS treated with DMSO showed a baseline level of PKA phosphorylation that was further increased by treatment with Fsk/IBMX (**Fig. 6D**). The prior *in vivo* experiments demonstrate phosphorylation of total cellular Tns3. Therefore, to determine whether the FA pool of Tns3 is also phosphorylated by PKA, Tns3 was immunoprecipitated from purified FACS proteins as well as from a comparable amount of whole cell extract and analyzed by immunoblotting. The amount of protein recovered from FACS preparations is limiting, leading to relatively poor quality of the blots. Nonetheless, Tns3 recovered from purified FACS fractions showed immunoreactivity with the phospho-PKA substrate antibody (**Fig 6E**). Taken together, the direct modification of Tns3 by PKA *in vitro*, the increased phosphorylation of Tns3 by Fsk and IBMX *in vivo*, and the presence of PKA-phosphorylated Tns3 in isolated FACS preparations strongly support the contention that the FA protein Tns3 is a new, direct substrate for PKA.

**Figure 6.**
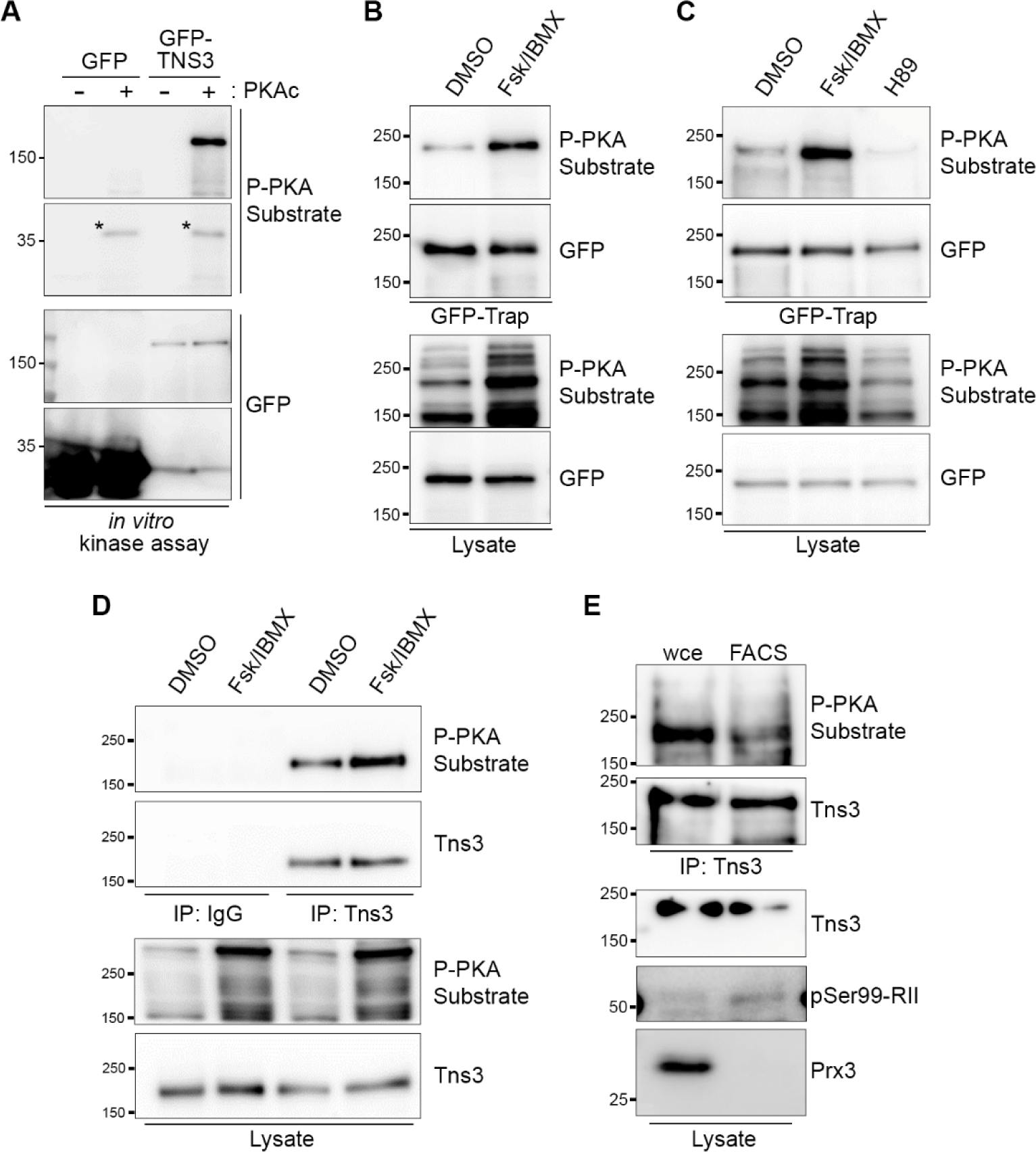
Tensin-3 (Tns3) is a direct physiological substrate for PKA. (**A**) HEK293T cells were transfected with GFP or GFP-Tensin3 (*GFP-TNS3*), lysed at 48 hr post transfection, and GFP-Trap pulldowns were incubated *in vitro* with ATP in the absence or presence of PKA catalytic subunits for 30 min. Reaction products were separated by SDS-PAGE and immunoblotted with anti-GFP or anti-phospho-PKA substrate (*P-PKA*) antibodies. (**B**) HEK293T cells were transfected with GFP-Tns3, serum-starved at 32 h post-transfection, and 16 h later treated with Fsk and IBMX (50 µM and 100 µM) or DMSO for 10 min. Cell lysates and GFP-Trap pulldowns were analyzed as described in (**A**). (**C**) HEK293T cells were transfected with GFP-Tns3 and treated 48 h post-transfection with DMSO, Fsk + IBMX (25 µM and 50 µM) or H89 (20 µM) in complete media for 10 min. Lysates and GFP-trap pulldowns were immunoblotted as described in (**A**). The positions of molecular weight markers, in kDa, are shown. (**D**) U2OS were serum-starved for 16 h then treated for 10 min with either DMSO (1 µl/ml) or Fsk/IBMX (25 µM/ 50 µM) in serum-free media. Lysates were immunoprecipitated with either rabbit IgG or anti-Tns3 antibody then immunoblotted as indicated. (**E**) Tns3 immunoprecipitates from 8 µg of whole cell extract or purified FACS proteins, along with 2 µg of each lysate, were analyzed by immunoblotting with the indicated antibodies. Positions of molecular weight markers are indicated throughout.

## DISCUSSION

Cell adhesion and migration are complex processes that are regulated by distinct and dynamic signaling events occurring within discrete subcellular regions and structures. FAs are multi-component junctions between ECM-bound integrins and the actin cytoskeleton whose complex constituents provide structural integrity as well as the capacity for dynamic turnover and for generating signals that convey and control the state of cell adhesion (3,4). PKA is a ubiquitous and promiscuous kinase with pleiotropic effects on cell adhesion and migration (17,21,27,32,33,35,36,38–42). Importantly, PKA activity is physically and functionally focused by subcellular localization through AKAPs, which is critical for establishing specificity and fidelity of PKA function (14–16). Here, we have shown that PKA is present and active within FAs and have identified several new putative FA substrates for PKA, including Tns3, a regulator of adhesion, migration, and cancer cell invasion and of signaling events associated with those cellular processes. These observations establish an important new niche for PKA signaling that is likely to play an important role in its effects on adhesion and migration.

We’ve demonstrated that inhibition of PKA affects FA morphology and distribution during fibroblast cell adhesion and spreading. Specifically, FAs are larger, longer, and more clustered, and show more peripheral distribution in PKA-inhibited cells. This phenotype resembles those described both in FAK-null keratinocytes (72) and FAK-null MEFs reconstituted with an active FAK-Src chimera (73), and is quite similar to a recently reported lack of FA ‘splitting’ during maturation (50). The fact that inhibition of PKA roughly phenocopies loss of FAK function, with respect to FA morphology, is particularly notable given the recent observation that FAK activity is required for localized activation of PKA in the leading edge of migrating cells (74). However, it is important to point out that the FA effects of PKA inhibition reported here are the result of global inhibition, *i.e.* inhibition of total cellular PKA activity. Given the current demonstration of PKA activity within FAs, it will be important – although technically challenging – to specifically inhibit or disrupt the FA pool of PKA and determine the consequences for adhesion and migration. That goal may also be realized or facilitated by investigating the effects of phosphomimetic and/or phosphoresistant mutants of various FA substrates for PKA.

To that end, the current work has identified a number of new potential FA substrates for PKA. We note that the approach used in this report is quite selective, as important, but more loosely-associated structural and regulatory proteins may not be retained throughout the highly-reductive FACS preparation procedure are thus not present for analysis as potential substrates. Furthermore, proteins that do survive fractionation but that are already highly phosphorylated at PKA-targeted sites will also not be efficiently modified or identified. This population could theoretically be recovered by treatment of lysates with alkaline phosphatase or the like prior to incubation with exogenous kinase. With these considerations in mind, the list of putative FACS substrates for PKA generated here, while substantial and intriguing, should not be considered exhaustive. Nonetheless, the number and variety of hits reported here not only adds to our understanding of PKA-mediated regulation of cell adhesion but significantly underscores and expands the utility and scope of kinase-catalyzed biotinylation as an effective chemical biology method for substrate identification (57,58,75). Specifically, the current work has demonstrated the efficacy of kinase-catalyzed biotinylation for substrate discovery using a much smaller amount of starting material than previously described. This lends itself to application of the technique to investigate other, specialized subcellular fractions of interest that may be difficult to isolate in large amounts.

Among the new putative substrates for PKA in FACS, this work further validated Tns3 as a *bona fide* physiological PKA substrate. While biochemical data presented here support the notion that Tns3 recovered from isolated FACS fraction is indeed phosphorylated by PKA, they do not establish that it is *specifically* or *exclusively* the pool of Tns3 within FAs that is phosphorylated by PKA. This would require either quantitative phospho-analysis of Tns3 isolated from purified FACS preparations vs cytosolic or immunofluorescence analysis with a phospho-specific antibody recognizing PKA site(s) on Tns3. This in turn requires identification of the PKA phosphorylation site(s) on Tns3, confirmation of these sites through *in vivo* testing, and generation and validation of phospho-specific antibodies. Clearly, considerably more time, effort, and expense are required – and planned – in order to determine whether it is the newly-identified FA pool of PKA that is modifying Tns3.These limitations notwithstanding, direct phosphorylation of Tns3 by PKA remains intriguing, given that one aspect of the FA phenotype observed in PKA-inhibited cells is a decrease in adhesive complexes in the cell center (**Fig. 1**), a location often associated with formation of fibrillar adhesions. Although we have not determined the site (s) of phosphorylation, human Tns3 contains several sites that score highly on *in silico* algorithms for predicting PKA phosphorylation sites (*e.g.* NetPhos3.1 (76) (http://services.healthtech.dtu.dk/services/NetPhos-3.1; **Fig. 5C**); pkaPS (http://mendel.imp.ac.at/pkaPS)). None of the highest-scoring predicted sites, however, are located in or near functionally well-characterized regions of Tns3 such as the N-terminal PTP or C2 or C-terminal SH2 or PTB domains, but occur instead in regions with very low confidence scores in terms of structure as predicted by AlphaFold (https://www.uniprot.org/uniprotkb/Q68CZ2/entry#structure), so their locations do not offer any obvious hints as to potential mechanistic consequences. However, many of the sites are conserved across Tns1-3 and, notably, both Tns2 and Tns3 were identified as hits with decreased phoshopeptide abundance in a CRISPR-based systems-level approach using CRISPR to knockout PKA expression and identify substrates in epithelial cells (77). This raises the possibility that the tensins represent a significant family of new PKA targets in cell adhesion. Clearly, it will be of considerable importance to identify the PKA-phosphorylated site (s) in Tns3, to determine whether other isoforms are similarly modified, and to characterize the role that this new mode of modification plays in regulating tensin biology.

The work reported here demonstrates that PKA RII subunits are associated with adhesion complexes, a finding corroborated by meta-analysis of published adhesion proteomes (1,51,53), and that FA contain subdomains of dynamic PKA activity. It would be inaccurate, however, to describe PKA as a ‘FA protein’. Unlike any ‘true’ FA protein, PKA is not exclusive to – or even significantly enriched in – FAs. However, a vast literature has established that PKA is it is not the bulk amount or relative abundance of PKA in a given subcellular niche that determines its importance. For example, only a miniscule fraction of total PKA is associated with AMPA receptors, yet this small fraction nonetheless plays a crucial role in neuronal plasticity (78–80). Many more examples exists, as PKA has hundreds of substrates throughout the cell and regulates diverse cellular processes ranging from regulation of ion channels at the plasma membrane to regulation of transcriptional and genomic events in the nucleus and myriad processes and locations in between (12,14,15,17,19,81,82). In essentially all of these cases, fidelity in the regulation of those processes is dependent upon anchoring of PKA to the regions or structures involved. Thus, the potential importance of PKA to FA function can be inferred from its existence in FA, as demonstrated here.

In that regard, another intriguing aspect of the current work is the demonstration of dynamic PKA activity within sub-domains of individual FAs. A significant literature has established that, rather than being uniform and homogenous structures, individual FAs are molecularly and spatially diverse, exhibiting regional differences in composition, distribution, cytoskeletal linkage, force transduction, and dynamics (83,84). For example, both nascent and mature, stable adhesions exhibit both stable and dynamic (*i.e.* low- and high-exchange) fractions of many if not most FA components, with turnover properties and rates that vary as a function of location with individual adhesions (83–86). Intriguingly, there is evidence to suggest that this variability correlates with – and may result from – differential phosphorylation of various FA components (2,84,87–90). The diversity of FA exists laterally (*i.e.* parallel to the plasma membrane) as well as in Z-dimension (84,91,92) and its complexity is further multiplied by cell-wide variability arising from the adhesion age (*i.e.* nascent *vs* stable) and position (*e.g.* lamellar vs central vs trailing edge) (84–86,92) and their regulation during migration (93). All of these considerations, plus the number of known and new potential PKA substrates in FAs, give rise to the intriguing possibility that the localized patterns of PKA may play a role in such sub-FA dynamics. Given the numerous variables and potential targets involved and the spatiotemporal resolution required to make correlative or causal connections, investigation of this possibility will require considerable, focused effort. Nonetheless, this demonstration of a discrete FA-associated pool of PKA activity raises many important questions. Principal among these are: (1) what anchors PKA within FAs? (2) how is PKA activity controlled in these structures? And (3) what are the targets for this activity and the consequences of their modification? Efforts to further address these questions and to expand our understanding of localized PKA signaling and the regulation of cell adhesion and migration are currently underway.

## EXPERIMENTAL PROCEDURES

### Reagents and antibodies

Rp-8-CPT-cAMPS (8- (4-chlorophenylthio)adenosine-3’,5’-cyclic monophosphorothioate, Rp-isomer) was purchased from either Biolog and Cayman Chemical with no notable differences in efficacy or potency. Reagents for immunofluorescence and immunoblotting were as follows: rabbit anti-paxillin (Y113; Abcam); mouse anti-vinculin (hVin-1; Sigma); rabbit anti-PKA RIIα (sc909; Santa Cruz Biotechnology); mouse anti-talin (8d4; Sigma); PKA-Cα (610980; BD biosciences); Prx3 (PF-PA0255; Abfrontier); rabbit monoclonal anti– phospho-RII (pS99 clone E151, ab32390; Abcam); mouse anti-tubulin (DM1a; Sigma); anti-GAPDH (14C10; Cell Signaling Technologies); mouse anti-actin (AC-40; Sigma); mouse anti-GFP (GF28R; Thermo Fisher); rabbit anti-phospho-PKA substrate (100G7E; Cell Signaling Technologies); and anti-Tns3 antibodies (PA5-63112; Invitrogen and HPA055338; Millipore-Sigma). Secondary reagents were purchased from Jackson ImmunoResearch (Alexa594- and Alexa647-conjugated donkey anti-rabbit and Alexa488-conjugated donkey anti-mouse) and Invitrogen (Alexa488-phalloidin; DAPI (4′,6-diamidino-2-phenylindole)). GFP-Trap® Magnetic Agarose was from ProteinTech. Reagents for synthesis of ATP-biotin were as described elsewhere (57). Unless otherwise noted, other common chemicals and reagents were from Sigma or Fisher.

### Plasmids and cloning

The plasmid encoding mCherry-paxillin was made by substituting mCherry for EGFP in pEGFP-N1-paxillin (a gift from Dr. Chris Turner (SUNY-Upstate)). The FA-targeted PKA biosensor PaxRAKS (paxillin-linked ratio-metric A-kinase sensor) was constructed using GAKdY (55), a single-fluorophore PKA biosensor which utilizes a PKA subtrate domain and FHA-derived phosphotheonine-binding domain to control the brightness of a conformationally-sensitive GFP variant. Specifically, the BamHI-NotI fragment from GAKdY (55) was ligated into the BamHI-NotI backbone of tdEOS-paxillin-22 (plasmid #57653; Addgene), replacing the tdEOS sequence to produce plasmid paxillin-GAKdy. The mCherry sequence from pmCherry-C1 (Clontech) was amplified to introduce flanking NheI sites using the following primers:

TTATGTAGGCTAGCGCTACCGGTCGCCACCATGGTGAGCA (fwd); AGCTGCTAGCTCGAGATCTGAGTCCGGACTTGTACAGCTCGTCCA (rev).

The NheI-digested PCR product was ligated into NheI-digested paxillin-GAKdY, fusing mCherry in-frame with the 5’ end of the paxillin-GAKdy sequence, producing mCherry-paxillin-GAKdY, which we refer to as PaxRAKS. The phospho-resistant control construct, in which the phosphoacceptor Thr residue in the PKA consensus sequence (RRA<u>T</u>) of GAKdY is replaced by Ala, was generated from PaxRAKS using site-directed mutagenesis (QuikChange II; Agilent). GFP-Tensin3 (catalog #105299) was purchased from Addgene.

### Cell culture and transfection

REF52 (a rat embryo fibroblast line kindly gifted by Dr. K. Burridge, University of North Carolina), HEK293T (human embryonic kidney epithelial cells; ATCC), U2OS (human osteosarcoma, ATCC), and B16F10 (mouse melanoma cells; ATCC) were routinely cultured in Dulbecco’s Modified Eagle’s Medium (MT10013CV; Corning/Fisher) containing 10% fetal bovine serum (A3160602; Gibco/Fisher). All cells were authenticated by STR profiling and tested for mycoplasma every six months. For transfection, 1 x 10^5^ cells were seeded in complete medium into 35 mm diameter tissue culture dishes and grown overnight, then transfected with 1.5 µg plasmid DNA using FuGENE 6 (Promega) as follows: 6 µl FuGENE 6 was diluted in 100 µl OptiMEM vortexed briefly and incubated at RT for 5 min; DNA was then added to the OptiMEM/FuGENE solution and the mixture briefly vortexed incubated at RT for 20 min; this solution was then added dropwise to the cells which were incubated for ∼48 h before further use.

### Immunofluorescence

Glass coverslips (22 mm; Electron Microscopy Services) were placed in a ceramic coverslip holder and sterilized by incubating in 2% HCl at 70°C for 30 minutes, washed in ddH_2_O twice for 10 minutes, incubated in a solution of 2% cuvette cleaning concentrate (Hellmanex III, Sigma) in ddH_2_O at 50°C for 30 minutes, and washed again in ddH_2_O twice for 10 minutes each. Coverslips were incubated in ddH_2_O at 90°C for 30 minutes, then in 70% ethanol at 70°C for 10 minutes and allowed to air-dry at 60°C overnight. Dry coverslips were stored in 70% ethanol until ready to use. Cleaned coverslips were removed from 70% ethanol and allowed to dry in a biosafety cabinet then were placed into the wells of a 6 well tissue culture plate and coated with 10 µg/ml fibronectin (BD Biosciences) in PBS (10.1 mM Sodium phosphate dibasic, 1.4 mM potassium phosphate dibasic, 137 mM sodium chloride, and 2.7 mM potassium chloride pH=7.4) at 37°C for 1 hour or overnight at 4°C. Coverslips were washed 3 times with PBS and prewarmed at 37°C before plating cells. Cells were trypsinized, resuspended in DMEM supplemented with 1% FBS and plated at ∼2000-5000 cells/cm^2^ for the specified time period before treatment and fixation. For fixation, culture medium was removed and cells were fixed in freshly prepared 4% formaldehyde in PBS for 15 minutes at RT, washed twice with PBS, permeabilized with PBS containing 0.25% triton X100 for 10 minutes at room temperature, and washed twice with PBS. For imaging of unroofed FA/cytoskeletal (FACS) samples, cells were fixed then unroofed using hydrodynamic shear from a dental water jet (Conair WaterpikTM WJ6RW), with a flow rate of ∼250 ml/min and a probe distance of ∼0.5 cm from the plating surface, rasterizing the probe across the surface for ∼10 sec. The coverslips were blocked in PBS containing 5% normal donkey serum (Jackson ImmunoResearch) for 1 h at RT or overnight at 4°C. The cells were then incubated with primary antibodies in a humidified staining chamber for 1 h at RT or overnight at 4°C. After three 5 min washes in PBS, cells were incubated with secondary antibodies and other reagents (phalloidin or DAPI) for 1 h at RT. After three 5 min washes in PBS, coverslips were mounted onto glass microscope slides (Fisher) using a small volume of PermaFluor mountant (Thermo Scientific) or in-house made mounting medium (prepared as described here: https://nic.med.harvard.edu/resources/media/). Epifluorescence images were captured through a 40x oil-immersion Plan Apo objective and fluorophore-specific filters (Chroma Technology Corp) on a Nikon Ti series inverted microscope equipped with a charged coupled device camera (Clara II, Andor Technologies) controlled by Nikon Elements software.

### Live-cell imaging

Cell spreading assays to assess focal adhesion morphology and evaluate traction forces in spreading cells were carried out identically prior to imaging. REF52 cells (untransfected or expressing indicated plasmids) were trypsinized, suspended in DMEM with 10% FBS, collected by centrifugation at 1200 X g at RT for 5 min. The cell pellet was resuspended in 2 mL Ringers buffer supplemented with 1% FBS. For pharmacological PKA inhibition, Rp-8-CPT-cAMPs (“Rp-cAMPs”, 75 µM) or 15 µl sterile H_2_O was added to cell suspension and gently mixed by trituration. Cell suspension was then plated onto prewarmed fibronectin coated 35 mm glass bottom imaging dishes (CellVis). Cells were returned to the tissue culture incubator and allowed to adhere for 20 minutes prior to imaging. Cells were maintained at 35-37°C using a NEVTEK ASI 400 Air Stream Incubator and a thin layer of light mineral was overlaid on the culture medium to prevent evaporation. For live-cell imaging of PaxRAKS, cells were imaged on a Nikon A1R confocal microscope with a beam splitter and dual GaAsP photomultiplier tube detectors, allowing simultaneous acquisition of both green- and red-fluorescence channels which, in turn, supports high acquisition frame rate and ratiometric two-color acquisition with no changes in sample morphology between channels. Lasers were powered at 2.5-2.8% with a scan speed of 15, and an HV/gain ratio = 45. Raw images were denoised using Nikon Elements.

### Focal adhesion analyses

Images of fixed REF52 and HDF cells taken using 40x oil immersion or 60x water immersion objectives were analyzed using a custom macro tool set created in FIJI (https://imagej.net/software/fiji/), which is freely available upon request. Briefly, images were imported into the program and underwent median filter processing (1.5) and background subtraction (rolling ball radius=50). The images were then thresholded to produce a binary image allowing isolation of FA and morphometric analysis of area and shape descriptors. In order to maintain some degree of automaticity while still considering cell-cell variation, the macro was written to produce several thresholded images in which the lower threshold was set anywhere from 9-15 times the mean of the background subtracted image. The most representative thresholded image was selected manually for analysis. Representative images were produced by thresholds across this spectrum regardless of timepoint, cell line, or treatment. FA area measurements and shape descriptors were calculated from the binary image using the *Analyze Particles* function in FIJI. Linearization or ‘unwrapping’ of cell images for determination of FA distance from center was performed using the *Polar Transformer* plugin of FIJI. Graphs and statistical analyses were produced using GraphPad Prism Software.

### Quantification of immunofluorescence colocalization

Two ImageJ plugins – *Coloc2* and *JACoP* (*J*ust *A*nother *Co*localization *P*lugin) – were used to quantify various colocalization parameters. Pearson’s correlation coefficient is a statistical method that characterizes the linear relationship between two variables. The most common method for quantifying colocalization in immunofluorescence images, it determines pixel-by-pixel covariance in the fluorescence signals between two images independently of signal intensity and background, providing a value of 1 in case of complete positive correlation, −1 for complete negative correlation (*i.e.* exclusion), and 0 for no correlation. Li’s intensity correlation coefficient (ICQ [PMID 15102922]) assesses the extent to which the signal *intensities* in two images covary while still removing bias for pixels of especially low or high intensity. ICQ values range from 0.5 to −0.5, where values closer to 0.5 indicating dependent staining, values near 0 indicating random staining, values near −0.5 for mutually-exclusive staining. Manders’ overlap coefficients quantify fractional overlap of two signals, ranging from 0 to 1. For the first fluorescence channel, the number of pixels (above a threshold value) that colocalize with thresholded pixels in the second channel is determined and divided by the total number of thresholded pixels in the first channel, defining M1 - the overlapping fraction for that channel. M2 conversely defines the proportion of thresholded pixels in the second channel that overlaps with first. Data were routinely tested for normal distribution using Prism software (GraphPad).

### Subcellular fractionation of FACS preparations

Focal adhesion/cytoskeleton (FACS) fractions were prepared using a modification of published methods (52). Briefly, cells were seeded into plates coated with 20 µg/ml fibronectin. After ∼21 h, cells (at ∼75% confluency) were fixed with dithiobis (succinimidyl propionate) (DSP, for immunofluorescence) or the reversible crosslinker dimethyl dithiobispropionimidate (DTBP, for biochemical analyses) at 37 °C for 15 min, followed by washing with complete PBS. The fixed cells were briefly incubated in ice-cold modified RIPA buffer (50 mM Tris-cl pH 7.5, 150 mM NaCl, 5 mM EDTA, 1% Igepal CA-630, 0.5% sodium deoxycholate) with protease inhibitor cocktail (Halt, ThermoFisher) and phosphatase inhibitors (10 mM NaF and 1 mM Na_3_VO_4_) for 2 min. Cell bodies and organelles were then removed by water shear (150 ml/s), and the remaining FACS complexes were either processed for immunofluorescence as described above or collected by scrapping in adhesion recovery solution (125 mM Tris pH 6.8, 1% SDS and 150mM DTT) and crosslinking was reversed by incubation at 37 °C for 30 min. The FACS proteins were acetone precipitated, resuspended in adhesion recovery buffer containing 25 mM DTT and stored at −20 °C until further use in immunoblotting or kinase-catalyzed biotinylation, as described below.

### Kinase-catalyzed biotinylation of FACS preparations

FACS protein preparations (5 µg for gel analysis, or 30 (trial 1) or 40 µg (trials 2-4) for LC-MS/MS) with or without added PKA (50 ng or 500 ng for gel analysis or 50 ng for LC-MS/MS, New England Biolabs) were combined with protein kinase buffer (PK buffer, New England Biolabs: 50 mM Tris-HCl, 10 mM MgCl_2_, 0.1 mM EDTA, 2 mM DTT, 0.01% Brij 35, pH=7.5). Reactions were initiated by adding ATP or ATP-biotin (2 mM) in a total 15 µL (gel analysis) or 20 µL (LC-MS/MS) volume and were incubated for 2 hours at 31 °C with 250 rpm mixing. After the 2 hr reaction time, samples were either immediately analyzed by gel analysis, as described below, or biotinylated proteins in the samples were enriched for LC-MS/MS analysis. Prior to enrichment, unreacted ATP-biotin was removed by diluting each reaction with phosphate binding buffer (PBB, 300 µL, 0.1 M phosphate pH=7.2, 0.15 M NaCl) and centrifugating in 3 kDa centriprep spin columns (Millipore, prewashed twice with 400 µL phosphate binding buffer) at 14,200 rcf for 30 min in 4 °C. After a second dilution and centrifugation step with the centriprep spin columns, as described above, samples were collected by centrifugation of the inverted centriprep column at 3200 rpm for 2 min at 4 °C. For biotinylated protein enrichment, each sample was diluted with PBS (300 µL) and combined with streptavidin agarose beads (GenScript, cat. No. L00353, 40 µL bead slurry, prewashed 3 times with 400 µL PBS). The bead/reaction mixture was incubated with end-over-end rotations for 1 hr at rt in spin columns (ThermoFisher, cat. No. 69705). After the 1 hr rotation, samples were spun down at 500 rcf for 1 min, and the collected beads were washed 5 times with PBS (400 µL) and 10 times with HPLC-grade water (400 µL). After washing, the biotinylated proteins were eluted with 2% SDS (100 µL) at 95 °C for 7 mins. The eluted proteins were concentrated using a SpeedVac (Thermo Scientific Savant SPD131DDA). Samples for both gel and LC-MS/MS analysis were denatured using Laemmli buffer (230 mM Tris-HCl, pH=6.8, 8% (w/v) SDS, 40 % (v/v) glycerol, 0.01 % (w/v) bromophenol blue, and 10% (v/v) β-mercaptoethanol) and heating at 95 °C for 5 min. For gel analysis, the denatured and biotinylated proteins were separated by 10% SDS-PAGE, electrotransferred onto PVDF membrane (Immobilon-P^SQ^, MilliporeSigma), and visualized using streptavidin-HRP (Cell Signaling) using an Alpha View-Fluorochem Q (ProteinSimple). For LC-MS/MS, the denatured proteins were desalted using 10% SDS-PAGE by running until the dye front was 1 cm from the well and stained with SyproRuby (Invitrogen) to visualize total proteins before sample preparation, as described below.

### LC-MS/MS sample preparation

HPLC grade solvents were used, and low binding Eppendorf tubes were washed once with water and once with acetonitrile. Proteins in each lane of the gel for LC-MS/MS analysis were excised, cut into small pieces (1 mm x 1 mm), and transferred to washed tubes. The gel pieces were incubated with ammonium bicarbonate (50 mM, 10 times the gel volume) for 10 minutes, the solution was discarded, and the gel pieces were incubated with acetonitrile until white and shrunk. The ammonium bicarbonate wash was repeated a second time, and the white and shrunk gel pieces were dried by SpeedVac. The dried gel pieces were incubated with TCEP (25 mM in 50 mM ammonium bicarbonate, 2 times the gel volume) at 37 °C for 20 minutes, and the TCEP solution was removed. The gel cubes were incubated with iodoacetamide (55 mM in 50 mM ammonium bicarbonate, 2 times the gel volume) for 1 hour with rocking, covered in foil, at RT, and the iodoacetamide solution was removed. The gel cubes were washed two times with a 1:1 mixture of 50 mM ammonium bicarbonate:acetonitrile with 15 minute incubation for each wash. Acetonitrile was then added to dehydrate the cubes, the acetonitrile was removed, and the samples were dried by SpeedVac. Gel pieces were rehydrated with trypsin solution (25 mM ammonium bicarbonate, 10% acetonitrile, 1:50 of trypsin:protein) and incubated at 37 °C overnight. The trypsin solution containing peptides was collected after centrifugation. Peptides in the gel pieces were extracted with extraction buffer (50% acetonitrile, 49.8% water, and 0.2% formic acid (ProteoChem)) with sonication for 15 minutes. The extraction solution was collected after centrifugation, combined with the trypsin solution, dried in a Speedvac, and stored at −20 °C until LC-MS/MS analysis at the Wayne State University and Karmanos Cancer Center Proteomics Core facility.

### LC-MS/MS analysis

Dried peptide digests were resuspended in 0.1% formic acid. Samples were separated by ultra-high-pressure reverse phase chromatography using an Acclaim PepMap RSLC column and an Easy nLC 1000 UHPLC system (Thermo). Peptides were analyzed with a Fusion mass spectrometer (Thermo) with a 120000 resolution orbitrap MS1 scan over the range of m/z 375− 1600, followed by ion trap resolution MS2 scans using an m/z 1.6 window and 30% normalized collision energy for HCD. Peak lists were generated with Proteome Discoverer (version 1.4; Thermo), and peptides scored using Mascot (version 2.4; Matrix Science) and Sequest (version 2.1). Label-free quantitation was performed using MaxQuant (version 1.6.3.4) software (59) (**Table S1A**). The digestion enzyme was set to trypsin/p with up to 2 missed cleavages. Carbamidomethylation of cysteine was set as a static modification. Oxidation of methionine and acetylation of the protein n-terminal were set as variable modifications. Identified unique and razor peptides were used for protein quantitation. The false discovery rate was set to 0.01 for identification of peptide spectral matches and proteins.

### Bioinformatics Analysis

Proteins with higher intensities in unlabeled samples (ATP+PKA) compared to samples treated with ATP-biotin and PKA (ATP-biotin+PKA) were removed. To identify candidate PKA substrates, enrichment values were calculated by dividing the protein intensities in the PKA-containing reactions (ATP-biotin+PKA) by the intensities of the same protein in the samples without added PKA (ATP-biotin-PKA). Fold enrichment values of either ≥1.5-or ≥6.0 in 2 out of 4 trials were used to generate hit lists (**Table S1, B and C**), with all remaining bioinformatic analysis done on the 53 hits in the ≥6.0-fold list (**Table S1C**). Functional analysis was performed using gene ontology (GO) with DAVID Bioinformatics Database (94). Previously identified physical protein– protein interactions among the 53 hits were mapped with Cytoscape 3.7.1 using the GeneMANIA 3.5.0 application (95,96). The presence of possible PKA motifs in each hit protein was assessed using NetPhos 3.1 using a probability cutoff of 0.5 (positive prediction) (76).

### Immunoblotting, immunoprecipitations, and pull-downs

For immunoblotting analysis, whole-cell extracts (WCE) were prepared lysing cells in ice-cold RIPA lysis buffer (150 mM NaCl, 1% Nonidet P-40, 0.5% sodium deoxycholate, 0.1% SDS, 50 mM Tris pH 7.4 with protease inhibitors (cOmplete; Millipore-Sigma) and phosphatase inhibitors (10 mM NaF and 1 mM Na_3_VO_4_)). For isolation of GFP-Tsn3, transfected cells were treated with solvent or pharmacological agents at 48 h post-transfection (either directly or after 16 h of serum starvation). Cells were incubated in either DMSO or Forskolin/IBMX (25 µM/ 50 µM) containing media for 10 min and lysed by RIPA buffer. The cell lysate was diluted in 10 mM Tris-Cl, pH 7.5, 150 mM NaCl, 0.5 mM EDTA and end-to-end rotated with 20 µl of GFP-Trap magnetic beads (ChromoTek) at 4° C for 2 h. The beads were then washed 4 times with cold dilution buffer containing 0.05% Igepal CA-630/NP-40. WCE, FACS fractions, or GFP-Trap pulldowns were boiled in Laemmli sample buffer then separated by SDS-PAGE, transferred to PVDF membranes, blocked with either 5% BSA or 5% NFDM in tris-buffered saline with 0.1 % Tween-20, and then immunoblotted with indicated primary and secondary antibodies. For control or Tns3 immunoprecipitations, 700 µg of RIPA lysates were incubated with 1 µg rabbit IgG (Jackson ImmunoResearch, # 011-000-003) or anti-Tns3 antibody (Invitrogen, PA5-63112) and rocked end-over-end at 4°C overnight, followed by incubation with 50 µl of protein A/G beads (Santa Cruz Biotechnology) at 4°C for 4 hr. The beads were then washed 4 times min with RIPA buffer and isolated proteins eluted by boiling in 2x Laemlli sample buffer, separated by SDS PAGE and immunoblotted with either anti-phospho-PKA substrate antibody (Cell Siginaling Technology, # 9621) or anti-Tns3 (Millipore Sigma, HPA055338).Ten µg of whole cell extract proteins were used for lysate controls. For immunoprecipitation of Tns3 from FACS fractions, U2OS cells were cross-linked, unroofed, and harvested as described above and Tns3 was immunoprecipitated from 8 µg of FACS protein or whole cell RIPA lysate and analyzed, along with 2 µg FACS or whole cell extract, by immunoblotting as indicated. Immunoblots were developed using enhanced chemiluminescence and signals captured by exposure to film or on a CCD-camera-based digital documentation system (BioRad).

### *In vitro* kinase assay

GFP- or GFP-Tensin-3 expressing cells were lysed in RIPA buffer (10 mM Tris/Cl pH 7.5, 150 mM NaCl, 0.5 mM EDTA, 0.1 % SDS, 1 % Triton™ X-100, 1 % deoxycholate), diluted 1:1 with dilution buffer (10 mM Tris/Cl pH 7.5, 150 mM NaCl, 0.5 mM EDTA; buffer recipes from the manufacturer (Chromotek)), and clarified by centrifugation at 17000 x g for 5 min at 4° C. Approximately 700 µg of each lysate were used for pulldowns by incubation with 25 µl GFP-Trap magnetic agarose (Chromotek) for 1 h at 4° C and washing four times with 10 mM Tris/Cl pH 7.5, 150 mM NaCl, 0.05 % IGEPAL CA-630, 0.5 mM EDTA) using a magnetic rack. After the final wash, the beads were resuspended in a small volume of wash buffer, divided into two new microfuge tubes, and incubated with 200 µM ATP with ot without 1 µl of recombinant PKA catalytic subunit (PKAc; New England Biolabs) in manufacturer-supplied reaction buffer at 30° C for 30min with agitation. Reactions were stopped by addition of an equal volume of 2X Laemmli sample buffer and boiling for 5 min, then products were run on duplicate SDS-PAGE gels (1/2 elute per lane) and immunoblotted with phospho-PKA substrate or GFP antibody.

## ACKNOWLEDGEMENTS

The authors thank John Patterson and Zoe Edmunds for technical assistance in the Howe Laboratory. This work was funded by NIH grants R01GM117490 and R01GM137611 (to AKH), R35GM131821 (to MKP), and P30ES020957, P30CA022453 and S10OD030484 (to Wayne State University and Karmanos Cancer Center Proteomics Core). Gene ontology analysis was performed in part by the Vermont Integrative Genomics Resource DNA Facility and supported by the University of Vermont Cancer Center, Lake Champlain Cancer Research Organization, and the UVM Larner College of Medicine.

## AUTHOR CONTRIBUTIONS

A.K.H. conceived and managed the study. M.K., A.J.S., H.N., M.M., R.J.B., A.A.H., and A.K.H. performed experiments and generated reagents. All authors analyzed data. M.K., A.J.S., M.M., R.J.B., A.A.H., MK.H.P., and A.K.H. generated and edited manuscript sections and figures. MK.H.P. and A.K.H. supervised the work and procured funding.

## DATA AVAILABILITY

All original data supporting the reported findings, as well as new reagents and software, are available from the corresponding author upon reasonable request. Macros and datasets are also available through Github (https://github.com/howelabuvm/Kang_JBC2024).

---

**Supporting Figure S1.**
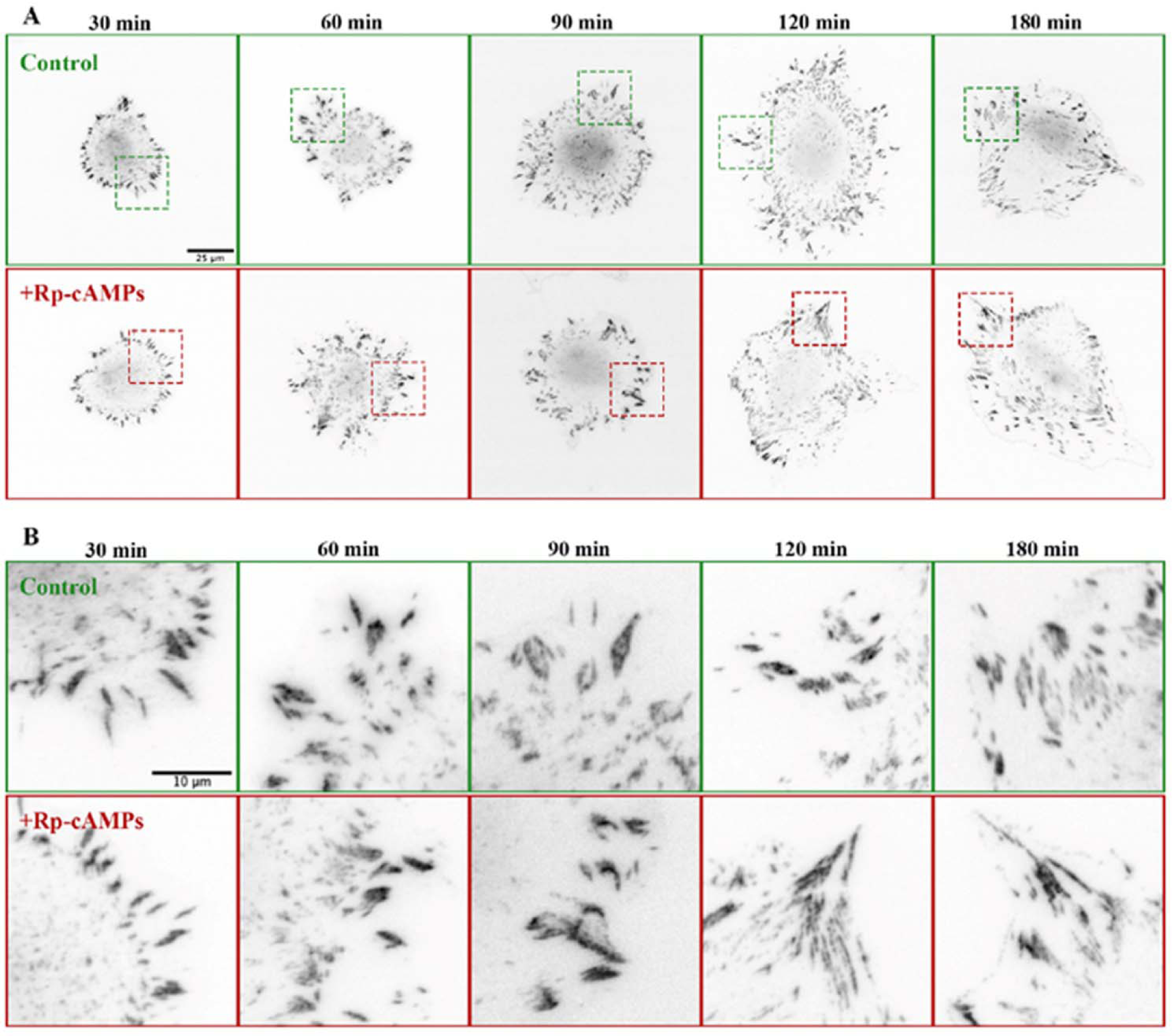
PKA inhibition changes FA morphology in spreading REF52 cells. (**A**) REF52 cells plated on fibronectin coated glass in the absence (control) and presence of the selective PKA inhibitor (Rp-8-CPT-cAMPS (*+Rp-cAMPS*); 50µM) were fixed 30, 60, 90, 120 and 180 minutes after plating and stained for paxillin. (**B**) Insets from cells highlighting morphological differences between groups upon treatment with Rp-cAMPs.

**Supporting Figure S2.**
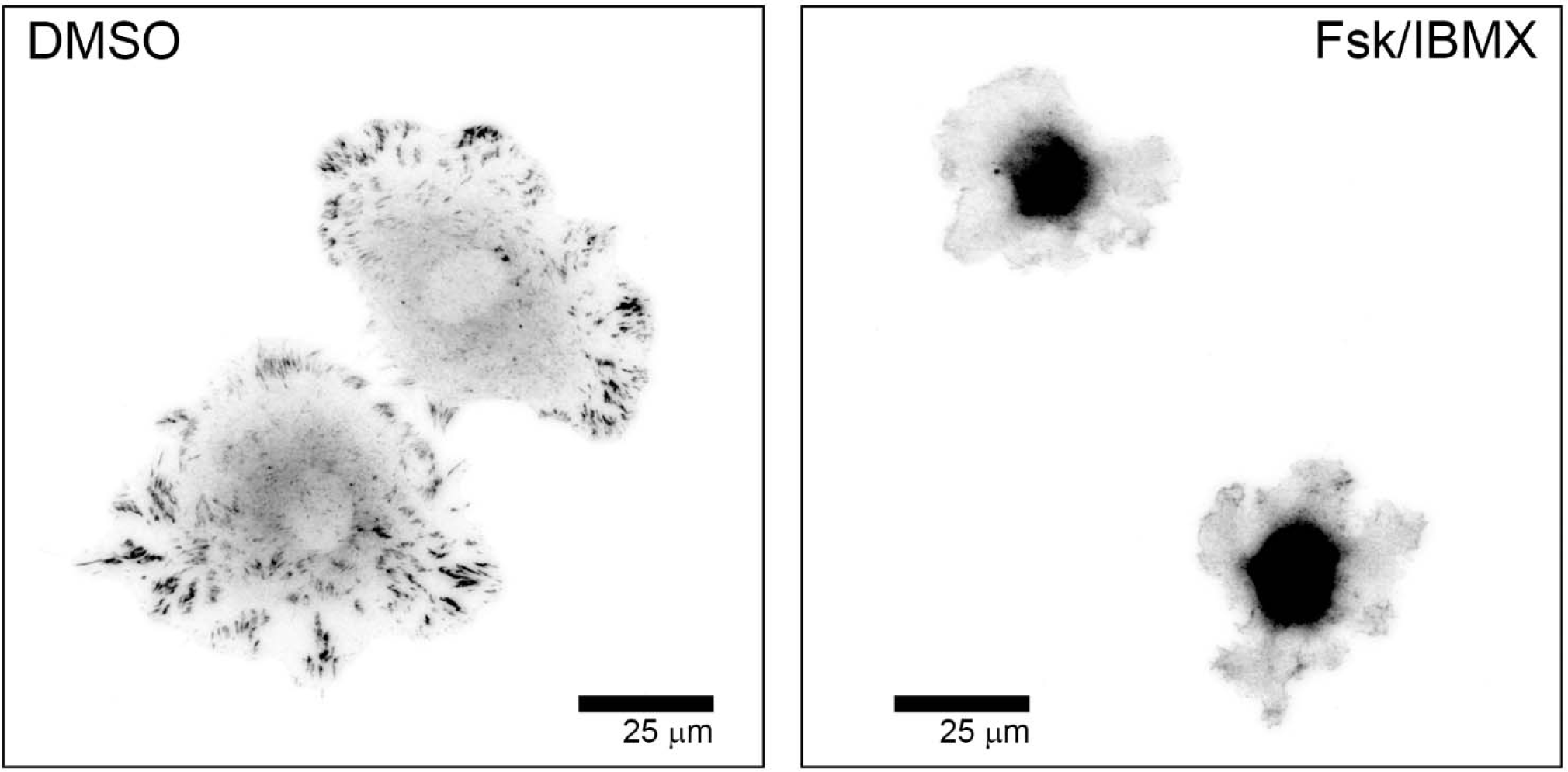
Addition of forskolin + IBMX inhibits cell spreading and FA formation. REF52 cells were plated on fibronectin-coated coverslips for 45 min in the presence of solvent (*DMSO*; 0.1% vol:vol) or a combination of forskolin and IBMX (*Fsk/IBMX*; 25 & 50 µM, respectively) to activate adenylyl cyclase and inhibit phosphodiesterase and fixed and stained for vinculin.

**Supporting Figure S3.**
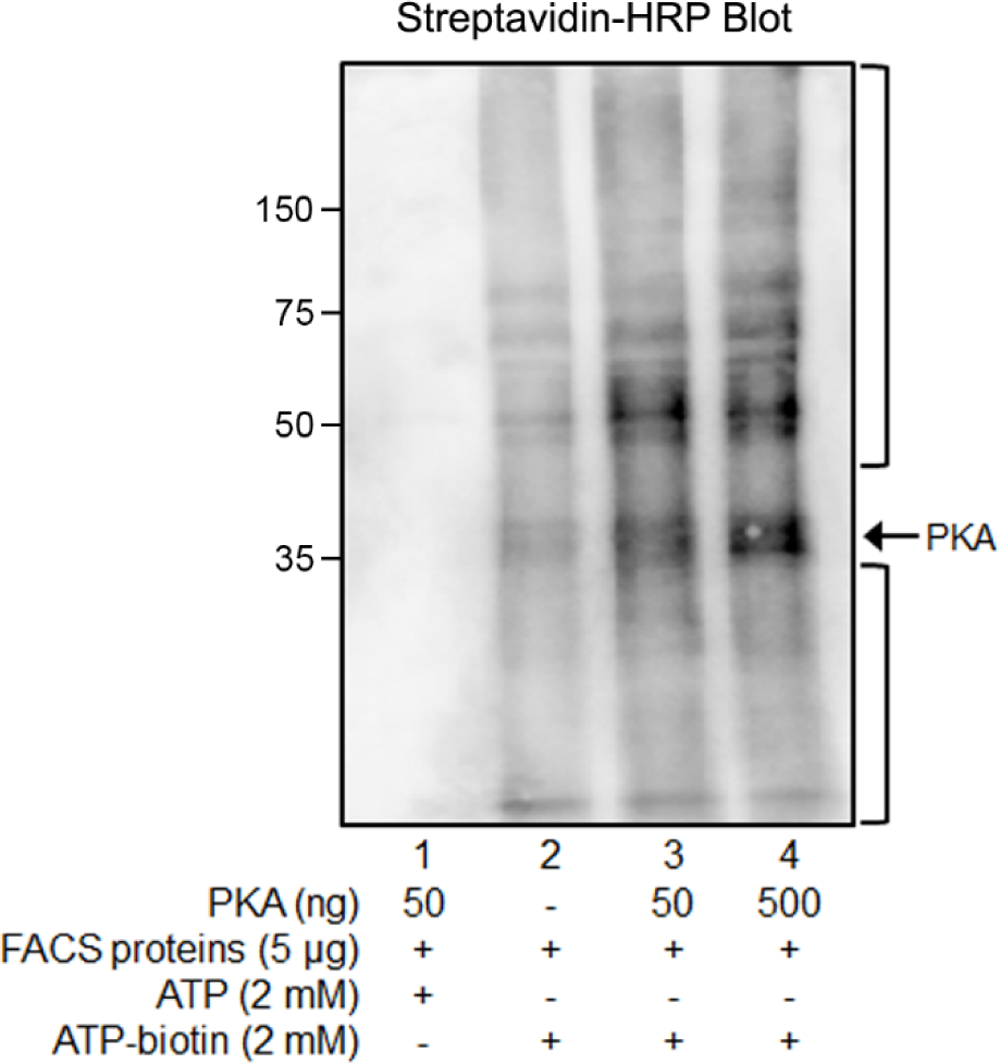
PKA-catalyzed biotinylation of isolated FACS proteins. Focal adhesion/ cytoskeleton (FACS) fractions were prepared from FN-adherent REF52 cells and processed as described in *Materials and Methods*. Kinase-dependent biotinylation was performed on FACS protein (5 µg) with either ATP (2mM, lane 1) or ATP-biotin (2 mM, lanes 2-4) in the absence (lane 2) or presence (lanes 1, 3, and 4) of exogenously added PKA (50 or 500 ng, as indicated) for 2 hours with 250 rpm mixing at 31 °C. The biotinylated proteins were separated by SDS-PAGE, transferred o PVDF membrane, and visualized by streptavidin-HRP. PKA is labeled with an arrow, potential substrates are indicated with brackets, and molecular weight markers (in kDa) are shown at left.

**Supporting Figure S4.**
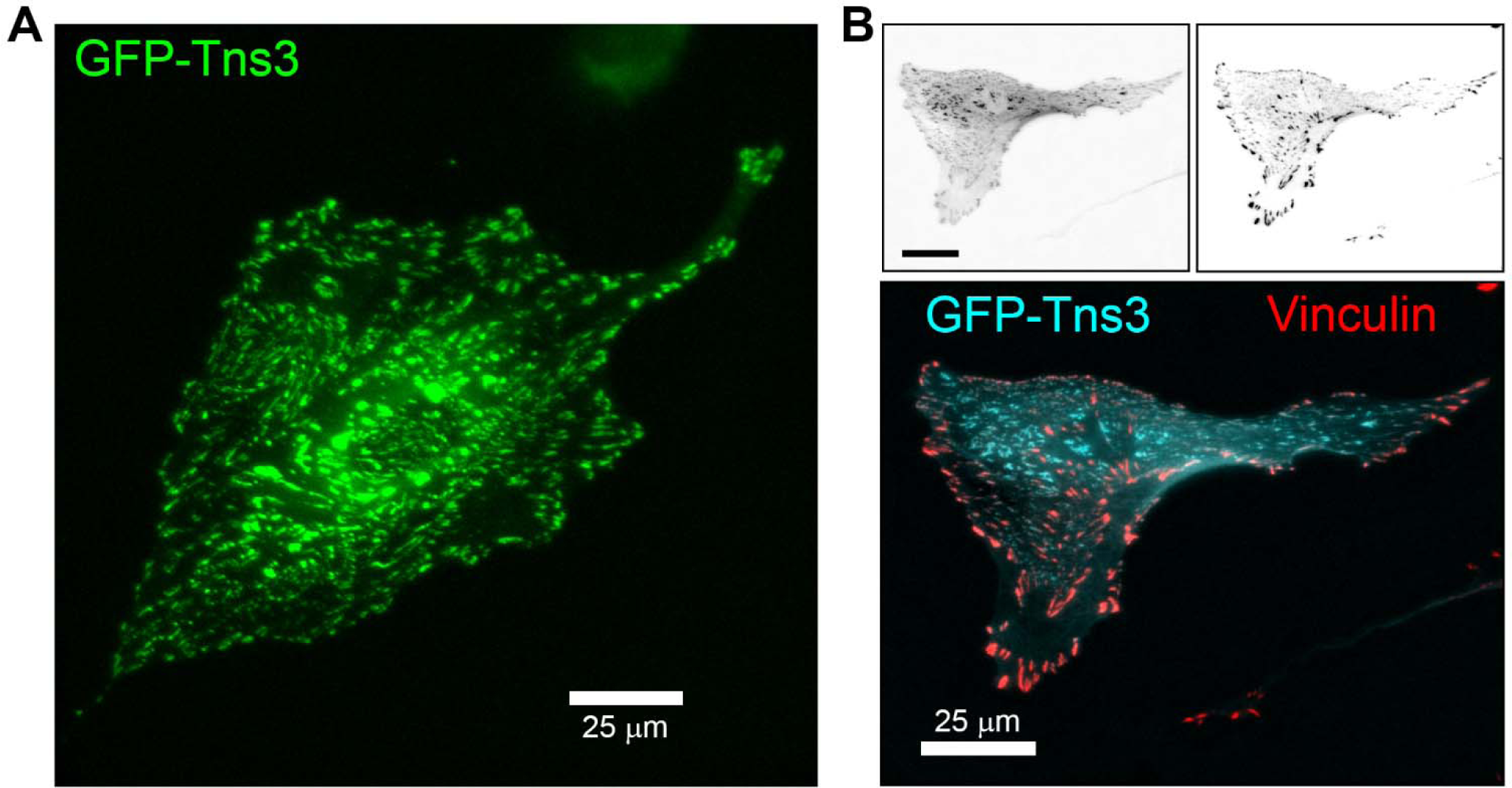
GFP-Tns3 localizes to focal and fibrillar adhesions. REF52 cells transfected with a plasmid encoding GFP-tagged Tns3 were plated onto FN-coated glass-bottom dishes and imaged by live-cell microscopy (**A**) or FN-coated coverslips and fixed, stained with antibodies against vinculin, and imaged by two-color fluorescence microscopy (**B**). Individual GFP-Tns3 and vinculin images are shown in inverted grayscale (*top left & right, respectively*) or pseudo-colored and overlaid (*bottom*).

## Notes

### Competing Interest Statement

The authors have declared no competing interest.

### Summary of Updates

New data showing the prominent localization of phospho-RII subunits to focal adhesions has been included (Fig. 3). Revisions have been made to (current) Figs. 1,2,4, and 6. Two new supplemental figures (Figs S2 and S4) have been added. Results, Discussion, and Experimental Methods sections have been updated to reflect additional data.

