## Supplementary figures and images for "Protein Kinase A is a Functional Component of Focal Adhesions"

### Supp Fig S1

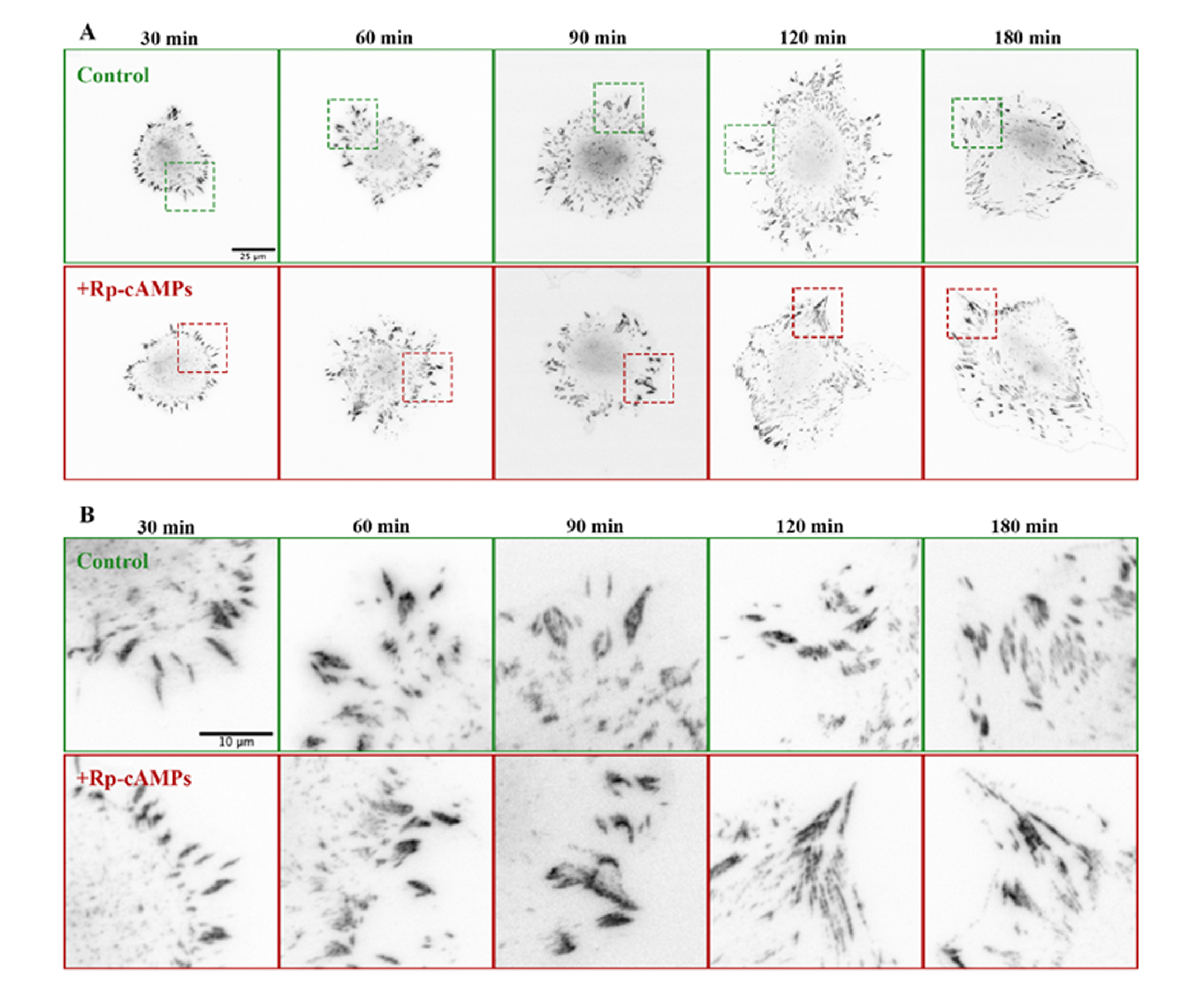

### Supp Fig S2

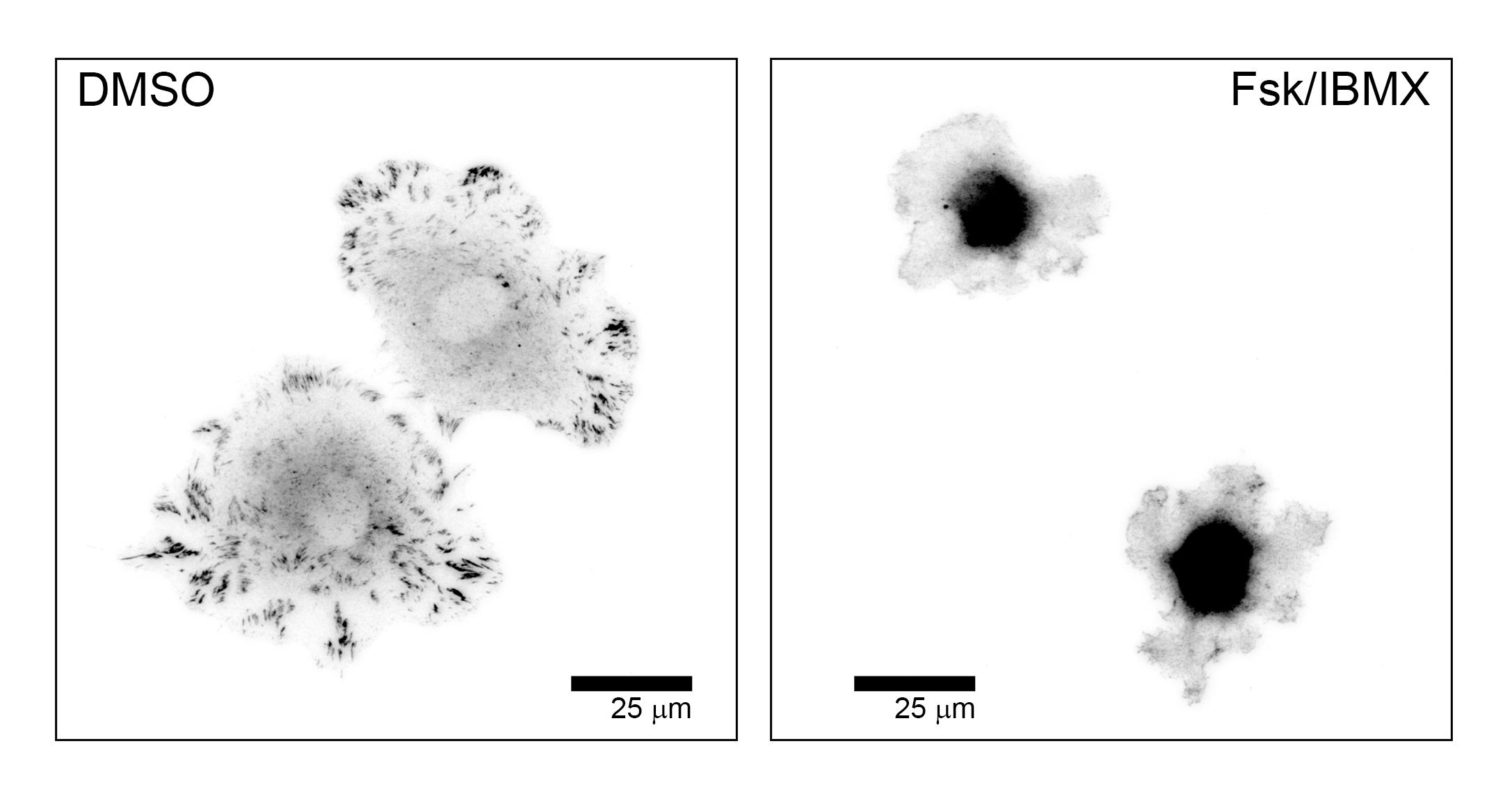

### Supp Fig S3

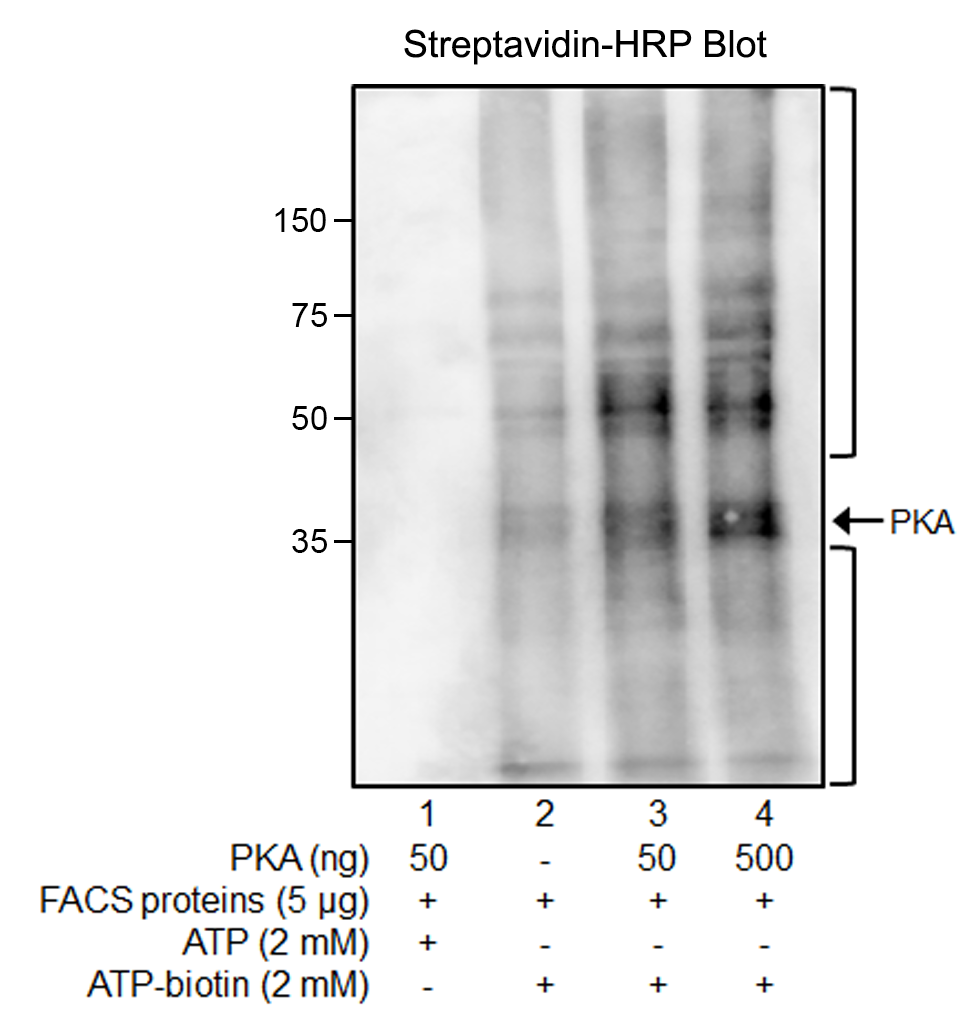

### Supp Fig S4

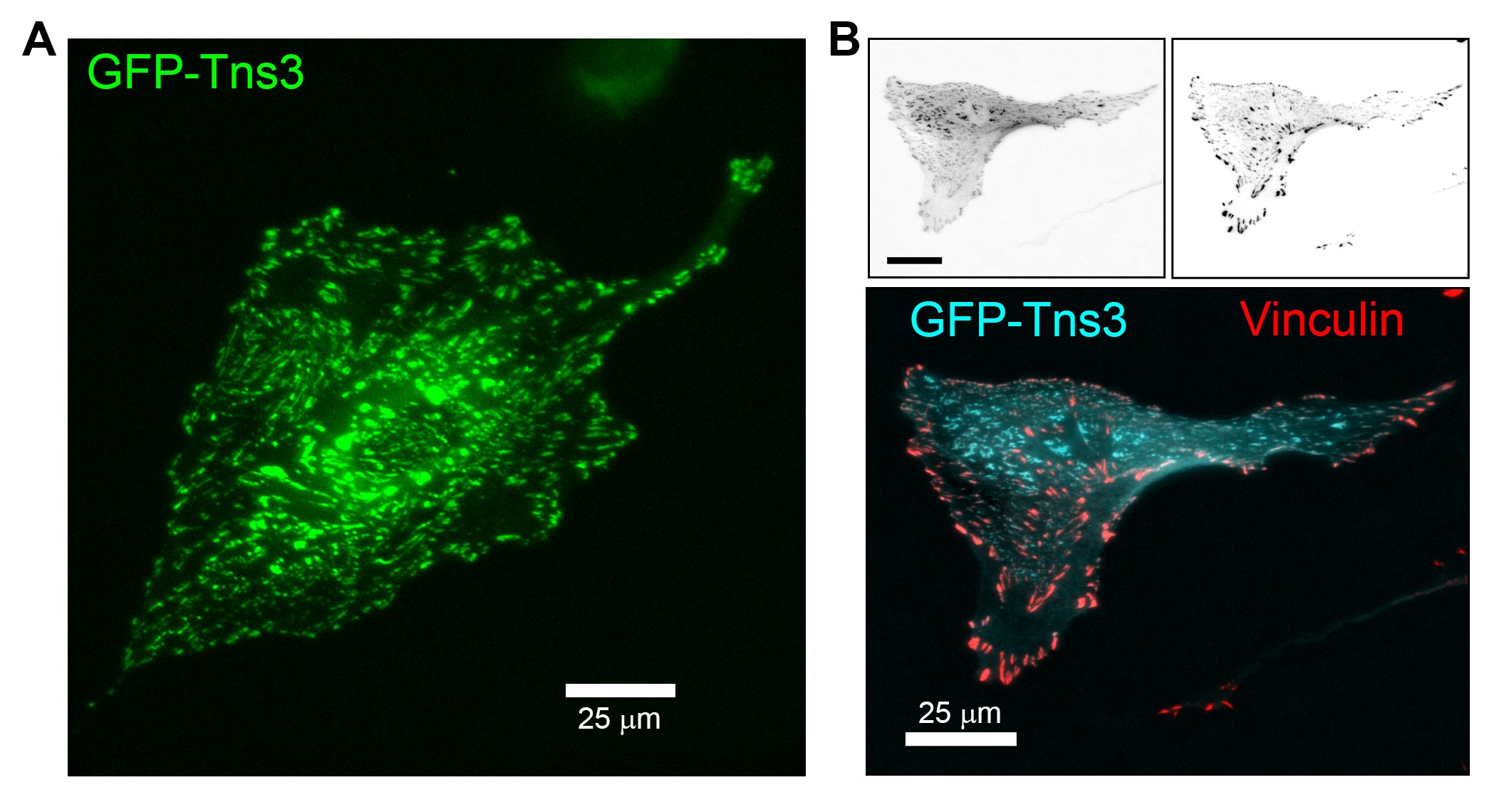
